# AlphaCRV: A Pipeline for Identifying Accurate Binder Topologies in Mass-Modeling with AlphaFold

**DOI:** 10.1101/2024.02.07.578780

**Authors:** Francisco J. Guzmán-Vega, Stefan T. Arold

**Affiliations:** Biological and Environmental Science and Engineering Division, Computational Bioscience Research Center, King Abdullah University of Science and Technology (KAUST), Thuwal 23955-6900, Kingdom of Saudi Arabia

**Keywords:** protein-protein interaction, clustering, scoring, high-throughput, rice, proteome

## Abstract

The speed and accuracy of deep learning-based structure prediction algorithms makes it now possible to perform in silico “pull-downs” to identify protein-protein interactions at a proteome-wide scale. However, existing scoring algorithms struggle to accurately identify correct interactions at such a large scale, resulting in an excessive number of false positives. Here, we introduce AlphaCRV, a Python package that helps identify correct interactors in a one-against-many AlphaFold screen by clustering, ranking, and visualizing conserved binding topologies, based on protein sequence and fold.

## Introduction

Protein-protein interactions are the foundation of most biological processes. With the advent of increasingly efficient and reliable computational methods, such as AlphaFold-Multimer (Evans et al. 2021) and other deep learning methods, rapid screening of protein-protein interactions has become feasible. This technological leap has paved the way for in silico identification and characterization of previously unknown complexes (Burke et al. 2023, Humphreys et al. 2021, O’Reilly et al. 2023). Recent enhancements in algorithmic efficiency have further accelerated modeling speed, enabling proteome-scale interaction predictions as a computational alternative to high-throughput experimental investigations (Ahdritz et al. 2022, Mirdita et al. 2022, Yu et al. 2023, Wallner 2023). However, the reliability of protein complex predictions lags behind that of monomeric structures (Bryant 2023), and a significant challenge remains in accurately identifying correct binders amidst thousands of predictions.

Active efforts are underway to develop computable quality scores for evaluating multimeric models (Bryant et al. 2022a, Bryant et al. 2022b, Malhotra et al. 2021, Zhao et al. 2023). Yet, at proteome levels, these scores (or their combinations) currently lack the selectivity needed to pinpoint the correct solutions, leading to a high number of false positives and rendering experimental verification impractical.

To address this challenge, we have developed AlphaCRV (an acronym for “AlphaFold Cluster, Rank, and Visualize”), a Python package designed to streamline the identification of true positives in a high-throughput AlphaFold screen of a bait protein against proteomes. The underlying principle of AlphaCRV is that a bait protein should bind similarly to homologous ligands in a one-against-many screen. Therefore, the more similar models with good quality scores and similar topology a given bait has, the less likely it is to be a spurious or “hallucinated” interaction. AlphaCRV clusters the top models based on sequence and fold to identify structurally similar binders, ranks them according to size and topological similarity, and generates files for PyMOL visualization.

AlphaCRV is a pipeline based on fast algorithms to cluster large sequence datasets in linear time (Steinegger and Söding 2018), and to cluster protein structures and identify remote homologies (Barrio-Hernandez et al. 2023). We show that AlphaCRV successfully identifies real binders in screens against the 43,000-protein rice proteome.

### Implementation

AlphaCRV is written in Python, with a command-line interface that allows the user to specify different parameters for the clustering process. It also allows the user to rank the resulting clusters and produce PyMOL sessions of the top clusters for easy visualization. To achieve this the program is separated into two stages that are described below.

#### Clustering stage

Starting from a directory with AlphaFold-Multimer models, which can be obtained easily with tools such as AlphaPulldown (Yu et al. 2023), the first stage of the pipeline retrieves the sequences and PDB models of the high-quality predictions and does both sequence and structure clustering to obtain a list of “joint clusters” (**Figure 1, upper panel**). The detailed steps are as follows:

**Figure 1.**
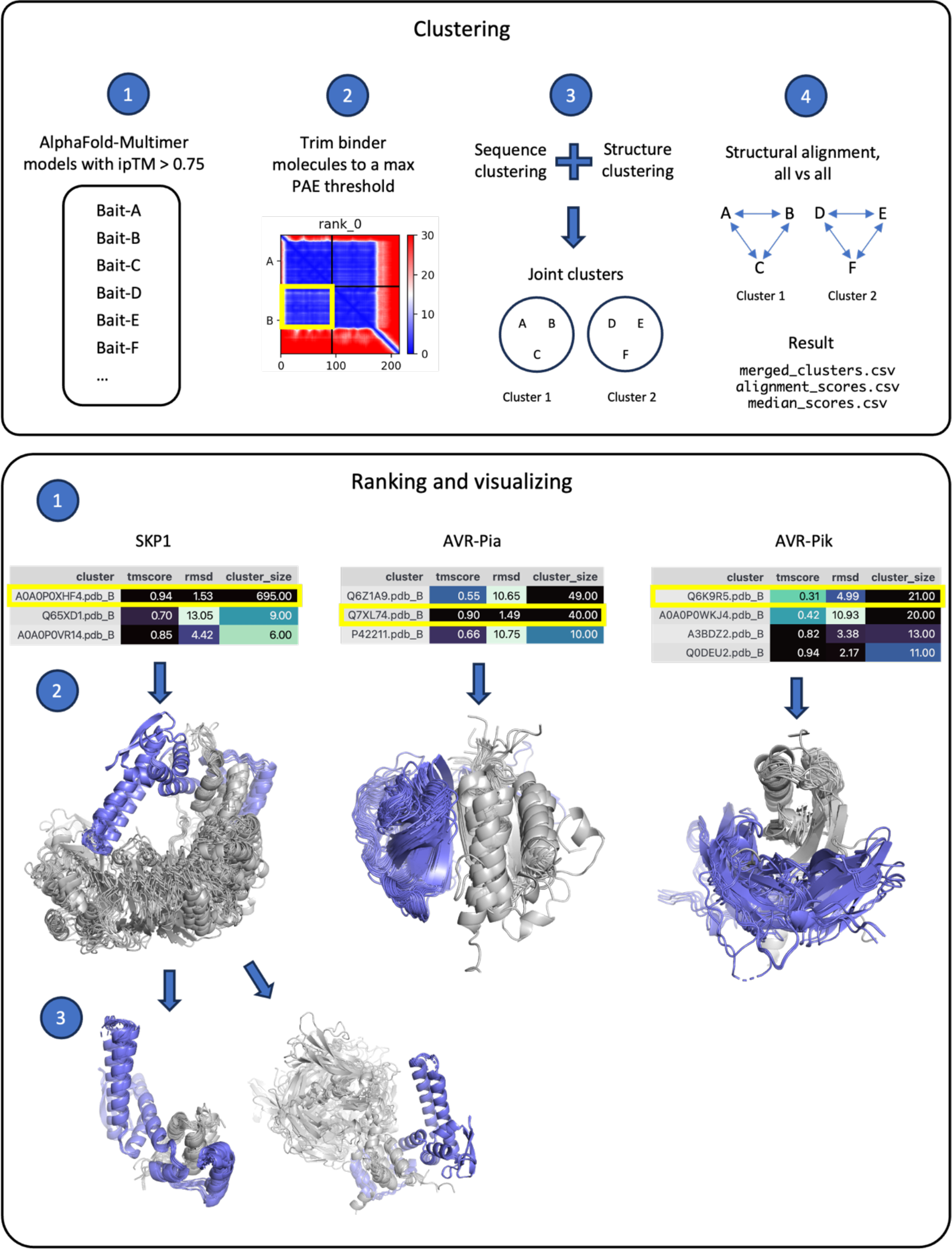
The AlphaCRV pipeline. The pipeline is divided in two stages: *Clustering* (upper panel), and *Ranking and Visualization* (lower panel). *Upper panel* (2) shows the PAE, colored from blue to red according to the estimated error in Å. X-axis shows residue numbers, and Y-axis shows segments for proteins A and B. The yellow box encircles the off-diagonal field indicating a possible interaction, which is used for trimming the ligands prior to clustering. *Lower panel (1)*: AlphaCRV was applied to a proteome-wide in-silico interaction screen of three proteins with different MSA depths: SKP1-like protein 20 (left), and AVR-Pia (center), AVR-Pik (right). The tables show the top-ranked clusters sorted by cluster size. The clusters containing the homologs of the true binder are highlighted in yellow. These clusters can be identified by the combination of a large cluster size and low RMSD of the cluster representative. *Lower panel (2)*: PyMOL session produced at the Visualization step. The bait proteins are colored in blue, and the (PAE-trimmed) rice interactors are gray. *Lower panel (3):* Subclusters that were identified within the top cluster of SKP1. The full analysis of these case studies is provided in the **Supplementary Note** and in the GitHub repository.

1. Obtain quality scores for the models and select the models with an interface predicted TM-score (ipTM) (Evans et al. 2021, Zhang and Skolnick 2004) equal or greater than the threshold set by the user (defaults to 0.75). This is done to only retain models with a high-quality interface prediction.
2. Use the Biskit Python library (Grünberg et al. 2007) to trim the PDB models and sequences based on a set predicted aligned error (PAE) (Evans et al. 2021, Jumper et al. 2021) threshold. This step only keeps the protein domains/regions of the binder that have a confident positioning with respect to the bait molecule.
3. Perform sequence clustering with mmseqs2 (Steinegger and Söding 2017, Steinegger and Söding 2018) on the trimmed sequences and structural clustering with foldseek (Barrio-Hernandez et al. 2023) on the trimmed structures. The resulting clusters are merged to create “joint clusters” that capture both sequence and structure relationships in the interacting molecules.
4. For each joint cluster, a pairwise all-vs-all structural alignment of the members is performed with US-align (Zhang et al. 2022) to find the best representative structure for each cluster. The best representative for each cluster is defined as the one with the lowest median RMSD when aligned to all other cluster members. The cluster representatives and their median RMSDs and TM-scores can then be used to rank the clusters.

The clustering stage produces as output csv files with a list of the clusters for each complex, and the alignment scores to rank the clusters. A detailed example of the analysis workflow is provided on the GitHub repository as Jupyter notebooks, which are also part of the **Supplementary Note**.

#### Ranking stage

The second stage of the pipeline ranks the clusters based on several user-defined criteria and produces PyMOL sessions for ease of visualization. Optionally, you can also request the program to further cluster the top clusters. This might be useful for very large clusters, when there are different binding modes predicted for the same topology of the binder or different interacting domains with a common binding domain. The detailed steps performed by the ranking stage are as follows (**Figure 1, bottom panel**):

1. Rank the clusters from the previous stage based on the cluster size, median TM-score, median RMSD and alignment coverage of the cluster representative.
2. Copy the PDB files of each of the top clusters to individual directories and make PyMOL sessions.
3. (Optional) Use foldseek once again to do structural clustering on the top clusters and save the results.

After this stage the program should produce a handful of clusters with promising binding topologies that can be easily inspected manually, from a starting list of potentially many thousands of models with similarly high interaction scores.

### Test case

As a test of our pipeline, we applied AlphaCRV to an interaction screen of three bait proteins with different multiple sequence alignment (MSA) depths against the rice proteome (*O. sativa subsp. Japonica)*. For this screen we used an in-house implementation of AlphaFold-Multimer v2.3 (Guzmán-Vega and Arold 2024) to produce 43,595 complexes for each bait: the SKP1-like protein 20 (UniProt ID Q651E8), an integral component of E3 ligase complexes, and two fungal effector proteins, AVR-Pia (B9WZW9), and AVR-Pik (C4B8B8), both of which have few or no known sequence homologues (**Figure 1**). For SKP1, which forms an extensive interface with its ligands, the ipTM and pDockQ scores ranked the true complex in 3^rd^ and 1^st^ place, respectively. However, the scoring differences to other solutions is too small to confidently identify the correct complex solely based on these scores (**Supplementary Table 1**). For AVR-Pia and AVR-Pik, ipTM ranked the correct solutions only as positions 440 and 237, whereas pDockQ ranked them as 168 and 140, respectively (**Supplementary Tables 2 and 3**). Conversely, AlphaCRV consistently identified the true binder and the correct topology as one of the top three largest clusters. In each case, the cluster with the true topology was the one with the best combination of large cluster size and low root mean square difference (RMSD). The ipTM and pDockQ scores alone were unable to identify the correct cluster for our test cases (**Supplementary Figure 1**).

We recommend first using AlphaCRV to identify the most likely correct topology. Subsequently, scores such as ipTM and pDockQ can be used to rank the models within each cluster, jointly with manual inspection of the complexes using the PyMOL files provided.

We note that AlphaCRV is designed to identify true positives from proteome-wide in silico pull-down screens, for example as a basis for future experimental investigations. False negatives are possible, for example if a bait uniquely binds to proteins with no homologs in the proteome.

## Availability

AlphaCRV is available for download in our GitHub repository (https://github.com/strubelab/AlphaCRV). It can be easily installed on any Linux machine through a conda environment. Detailed documentation and instructions for installation, usage, and examples are provided in the repository.

### Supplementary Data

Supplementary Tables and Figure, and the Supplementary Note with the detailed analysis of the examples are available online. The example datasets to run the case studies presented on this paper are available at https://zenodo.org/records/10470744.

## Acknowledgements

This research was supported by the King Abdullah University of Science and Technology (KAUST) through the baseline fund and the Award No. FCC/1/1976-33 and REI/1/4446-01 from the Office of Sponsored Research (OSR). For computer time, this research used the resources of the KAUST Supercomputing Laboratory (KSL). We thank N. Kathiresan and G. Wickham from the KSL for their support.

## Author contributions

FJGV and STA conceived the project. FGJV wrote the code. FJGV and STA wrote the manuscript.

## References

Ahdritz, Gustaf, Bouatta, Nazim, Kadyan, Sachin, Xia, Qinghui, Gerecke, William, O’Donnell Timothy J, et al., ‘OpenFold: Retraining AlphaFold2 Yields New Insights into Its Learning Mechanisms and Capacity for Generalization’, bioRxiv, 2022 <https://www.biorxiv.org/content/10.1101/2022.11.20.517210>

Barrio-Hernandez, Inigo, Yeo, Jingi, Jänes Jürgen, Mirdita, Milot, Gilchrist, Cameron L. M., Wein, Tanita, et al., ‘Clustering Predicted Structures at the Scale of the Known Protein Universe’, Nature, 622/7983 (2023), 637–45 <https://www.nature.com/articles/s41586-023-06510-w> [accessed 5 January 2024]

Bryant, Patrick, ‘Deep Learning for Protein Complex Structure Prediction’, Current Opinion in Structural Biology, 79 (2023), 102529 <https://linkinghub.elsevier.com/retrieve/pii/S0959440X23000039> [accessed 5 January 2024]

Bryant, Patrick, Pozzati, Gabriele, and Elofsson, Arne, ‘Improved Prediction of Protein-Protein Interactions Using AlphaFold2’, Nature Communications, 13/1 (2022a), 1265 <https://www.nature.com/articles/s41467-022-28865-w> [accessed 4 May 2023]

Bryant, Patrick, Pozzati, Gabriele, Zhu, Wensi, Shenoy, Aditi, Kundrotas, Petras, and Elofsson, Arne, ‘Predicting the Structure of Large Protein Complexes Using AlphaFold and Monte Carlo Tree Search’, Nature Communications, 13/1 (2022b), 6028 <https://www.nature.com/articles/s41467-022-33729-4> [accessed 7 November 2022]

Burke, David F., Bryant, Patrick, Barrio-Hernandez, Inigo, Memon, Danish, Pozzati, Gabriele, Shenoy, Aditi, et al., ‘Towards a Structurally Resolved Human Protein Interaction Network’, Nature Structural & Molecular Biology, 30/2 (2023), 216–25 <https://www.nature.com/articles/s41594-022-00910-8> [accessed 4 January 2024]

Evans, Richard, O’Neill, Michael, Pritzel, Alexander, Antropova, Natasha, Senior, Andrew, Green, Tim, et al., Protein Complex Prediction with AlphaFold-Multimer (4 October 2021) <http://biorxiv.org/lookup/doi/10.1101/2021.10.04.463034> [accessed 7 November 2022]

Grünberg, Raik, Nilges, Michael, and Leckner, Johan, ‘Biskit—A Software Platform for Structural Bioinformatics’, Bioinformatics, 23/6 (2007), 769–70 <https://academic.oup.com/bioinformatics/article/23/6/769/415036> [accessed 8 January 2024]

Guzmán-Vega Francisco J., Arold Stefan T. ’ strubelab/alphafold-ibex: Scaling up protein modeling with AlphaFold for SLURM-based HPC clusters, GitHub repository (2024) <https://github.com/strubelab/alphafold> DOI 10.5281/zenodo.10548184

Humphreys, Ian R., Pei, Jimin, Baek, Minkyung, Krishnakumar, Aditya, Anishchenko, Ivan, Ovchinnikov, Sergey, et al., ‘Computed Structures of Core Eukaryotic Protein Complexes’, Science, 374/6573 (2021), eabm4805 <https://www.science.org/doi/10.1126/science.abm4805> [accessed 4 January 2024]

Jumper, John, Evans, Richard, Pritzel, Alexander, Green, Tim, Figurnov, Michael, Ronneberger, Olaf, et al., ‘Highly Accurate Protein Structure Prediction with AlphaFold’, Nature, 596/7873 (2021), 583–89 <https://www.nature.com/articles/s41586-021-03819-2> [accessed 9 November 2022]

Malhotra, Sony, Joseph Agnel Praveen, Thiyagalingam, Jeyan, and Topf, Maya, ‘Assessment of Protein–Protein Interfaces in Cryo-EM Derived Assemblies’, Nature Communications, 12/1 (2021), 3399 <https://www.nature.com/articles/s41467-021-23692-x> [accessed 4 May 2023]

Mirdita, Milot, Schütze, Konstantin, Moriwaki, Yoshitaka, Heo, Lim, Ovchinnikov, Sergey, and Steinegger, Martin, ‘ColabFold: Making Protein Folding Accessible to All’, Nature Methods, 19/6 (2022), 679–82 <https://www.nature.com/articles/s41592-022-01488-1> [accessed 5 March 2023]

O’Reilly Francis J, Graziadei, Andrea, Forbrig, Christian, Bremenkamp, Rica, Charles, Kristine, Lenz, Swantje, et al., ‘Protein Complexes in Cells by AI-Assisted Structural Proteomics’, Molecular Systems Biology, 19/4 (2023), e11544 <https://www.embopress.org/doi/10.15252/msb.202311544> [accessed 4 January 2024]

Steinegger, Martin, and Söding, Johannes, ‘Clustering Huge Protein Sequence Sets in Linear Time’, Nature Communications, 9/1 (2018), 2542 <https://www.nature.com/articles/s41467-018-04964-5> [accessed 5 January 2024]

Steinegger, Martin, and Söding, Johannes, ‘MMseqs2 Enables Sensitive Protein Sequence Searching for the Analysis of Massive Data Sets’, Nature Biotechnology, 35/11 (2017), 1026–28 <https://www.nature.com/articles/nbt.3988> [accessed 5 January 2024]

Wallner, Björn, ‘AFsample: Improving Multimer Prediction with AlphaFold Using Massive Sampling’, ed. by Janet Kelso,Bioinformatics, 39/9 (2023), btad573 <https://academic.oup.com/bioinformatics/article/doi/10.1093/bioinformatics/btad573/7274860> [accessed 4 January 2024]

Yu, Dingquan, Chojnowski, Grzegorz, Rosenthal, Maria, and Kosinski, Jan, ‘AlphaPulldown—a Python Package for Protein– Protein Interaction Screens Using AlphaFold-Multimer’, ed. by Lenore Cowen, Bioinformatics, 39/1 (2023), btac749 <https://academic.oup.com/bioinformatics/article/doi/10.1093/bioinformatics/btac749/6839971> [accessed 29 April 2023]

Zhang, Chengxin, Shine, Morgan, Pyle Anna Marie, and Zhang, Yang, ‘US-Align: Universal Structure Alignments of Proteins, Nucleic Acids, and Macromolecular Complexes’, Nature Methods, 19/9 (2022), 1109–15 <https://www.nature.com/articles/s41592-022-01585-1> [accessed 5 January 2024]

Zhang, Yang, and Skolnick, Jeffrey, ‘Scoring Function for Automated Assessment of Protein Structure Template Quality’, Proteins: Structure, Function, and Bioinformatics, 57/4 (2004), 702–10 <https://onlinelibrary.wiley.com/doi/10.1002/prot.20264> [accessed 9 January 2024]

Zhao, Haiqing, Petrey, Donald, Murray, Diana, and Honig, Barry, ZIPPI: Proteome-Scale Sequence-Based Evaluation of Protein-Protein Interaction Models (25 May 2023) <http://biorxiv.org/lookup/doi/10.1101/2023.05.25.542344> [accessed 5 January 2024]

