## Supplementary Data for "AlphaCRV: A Pipeline for Identifying Accurate Binder Topologies in Mass-Modeling with AlphaFold"

| Rank by ipTM score | | | |
|---|---|---|---|
| rank | complex | ipTM | pDockQ |
| 1 | 8IF6-1_Q6Z1A9-1 | 0.9033 | 0.7391 |
| 2 | 8IF6-1_Q7XSL8-1 | 0.9029 | 0.7380 |
| 3 | 8IF6-1_Q5VMP0-1* | 0.8992 | 0.7401 |
| 8 | 8IF6-1_A0A0P0Y6A8-1 | 0.8976 | 0.7380 |
| 339 | 8IF6-1_Q8RZQ3-1 | 0.8671 | 0.7390 |
| 399 | 8IF6-1_A0A0N7KEW0-1 | 0.8625 | 0.7361 |
| 554 | 8IF6-1_Q69X07-1 | 0.8473 | 0.7288 |
| **Rank by pDockQ score** | | | |
| rank | complex | ipTM | pDockQ |
| 1 | 8IF6-1_Q5VMP0-1* | 0.8992 | 0.7401 |
| 2 | 8IF6-1_Q0E1Z5-1 | 0.8911 | 0.7398 |
| 3 | 8IF6-1_Q7XVM8-1 | 0.8888 | 0.7398 |
| 24 | 8IF6-1_Q8RZQ3-1 | 0.8671 | 0.7390 |
| 128 | 8IF6-1_A0A0P0Y6A8-1 | 0.8976 | 0.7380 |
| 363 | 8IF6-1_A0A0N7KEW0-1 | 0.8625 | 0.7361 |
| 675 | 8IF6-1_Q69X07-1 | 0.8473 | 0.7288 |

**Supplementary Table 1.** Ranking of complexes of SKP1 with true binder and homologous sequences of the true binder in rice. The top scored complexes overall that are not homologs are shaded grey. * Complex with full chain of true binder

| Rank by ipTM score | | | |
|---|---|---|---|
| rank | complex | ipTM | pDockQ |
| 1 | 6Q76-1_Q0IT61-1 | 0.9573 | 0.7082 |
| 2 | 6Q76-1_Q851F1-1 | 0.9550 | 0.7291 |
| 3 | 6Q76-1_Q653R0-1 | 0.9530 | 0.7294 |
| 159 | 6Q76-1_Q6EPT2-1 | 0.9138 | 0.7220 |
| 245 | 6Q76-1_Q6EPT4-1 | 0.9038 | 0.7172 |
| 343 | 6Q76-1_Q2QSQ7-1 | 0.8921 | 0.7202 |
| 388 | 6Q76-1_Q0J314-1 | 0.8863 | 0.7136 |
| 440 | 6Q76-1_6Q76A-1* | 0.8808 | 0.7250 |
| 488 | 6Q76-1_Q8S5W0-1 | 0.8757 | 0.7208 |
| 554 | 6Q76-1_Q6YY34-1 | 0.8673 | 0.7156 |
| 582 | 6Q76-1_Q7XJV3-1 | 0.8622 | 0.7171 |
| 592 | 6Q76-1_Q6YY33-1 | 0.8609 | 0.7127 |
| 624 | 6Q76-1_A0A0N7KFK3-1 | 0.8571 | 0.7128 |
| 652 | 6Q76-1_A0A0P0VKX7-1 | 0.8541 | 0.7129 |
| 724 | 6Q76-1_A0A0P0WB87-1 | 0.8431 | 0.6985 |
| 835 | 6Q76-1_Q6YY31-1 | 0.8251 | 0.7001 |
| 879 | 6Q76-1_Q0JCK8-1 | 0.8183 | 0.7016 |
| 948 | 6Q76-1_Q7XJV0-1 | 0.8064 | 0.6999 |
| 1438 | 6Q76-1_E9KPB5-1** | 0.6884 | 0.5947 |
| **Rank by pDockQ score** | | | |
| rank | complex | ipTM | pDockQ |
| 1 | 6Q76-1_A0A0P0X879-1 | 0.9117 | 0.7407 |
| 2 | 6Q76-1_Q5JMA6-1 | 0.9390 | 0.7365 |
| 3 | 6Q76-1_A0A0P0Y5V7-1 | 0.9004 | 0.7364 |
| 168 | 6Q76-1_6Q76A-1* | 0.8808 | 0.7250 |
| 242 | 6Q76-1_Q6EPT2-1 | 0.9138 | 0.7220 |
| 282 | 6Q76-1_Q8S5W0-1 | 0.8757 | 0.7208 |
| 292 | 6Q76-1_Q2QSQ7-1 | 0.8921 | 0.7202 |
| 364 | 6Q76-1_Q6EPT4-1 | 0.9038 | 0.7172 |
| 370 | 6Q76-1_Q7XJV3-1 | 0.8622 | 0.7171 |
| 412 | 6Q76-1_Q6YY34-1 | 0.8673 | 0.7156 |
| 474 | 6Q76-1_Q0J314-1 | 0.8863 | 0.7136 |
| 497 | 6Q76-1_A0A0P0VKX7-1 | 0.8541 | 0.7129 |
| 498 | 6Q76-1_A0A0N7KFK3-1 | 0.8571 | 0.7128 |
| 502 | 6Q76-1_Q6YY33-1 | 0.8609 | 0.7127 |

| | | | |
|---|---|---|---|
| 729 | 6Q76-1_Q0JCK8-1 | 0.8183 | 0.7016 |
| 749 | 6Q76-1_Q6YY31-1 | 0.8251 | 0.7001 |
| 753 | 6Q76-1_Q7XJV0-1 | 0.8064 | 0.6999 |
| 772 | 6Q76-1_A0A0P0WB87-1 | 0.8431 | 0.6985 |
| 1166 | 6Q76-1_E9KPB5-1** | 0.6884 | 0.5947 |

**Supplementary Table 2.** Ranking of complexes of AVR-Pia with true binder and homologous sequences of the true binder in rice. The top scored complexes overall (not homologs) are shaded grey. * Complex with true binder trimmed to only the portion in PDB 6Q76
** Complex with full chain of true binder

| Rank by ipTM score | | | |
|---|---|---|---|
| rank | complex | ipTM | pDockQ |
| 1 | 6R8K-1_A0A0P0Y5A4-1 | 0.9544 | 0.7326 |
| 2 | 6R8K-1_A0A0P0WTA7-1 | 0.9367 | 0.7401 |
| 3 | 6R8K-1_A0A0P0YB11-1 | 0.9165 | 0.7314 |
| 114 | 6R8K-1_Q7XJV3-1 | 0.8290 | 0.7321 |
| 211 | 6R8K-1_Q6EPT2-1 | 0.8069 | 0.7239 |
| 237 | 6R8K-1_6R8KA-1* | 0.8007 | 0.7268 |
| 240 | 6R8K-1_A0A0N7KFK3-1 | 0.8004 | 0.7330 |
| 315 | 6R8K-1_Q6YY31-1 | 0.7850 | 0.7303 |
| 428 | 6R8K-1_Q6EPT4-1 | 0.7668 | 0.7179 |
| 486 | 6R8K-1_Q0JCK8-1 | 0.7562 | 0.7255 |
| 3419 | 6R8K-1_D5L9G5-1** | 0.3917 | 0.3387 |
| **Rank by pDockQ score** | | | |
| rank | complex | ipTM | pDockQ |
| 1 | 6R8K-1_A0A0P0WTA7-1 | 0.9367 | 0.7401 |
| 2 | 6R8K-1_G9LZD7-1 | 0.8740 | 0.7393 |
| 3 | 6R8K-1_Q5NAK8-1 | 0.8294 | 0.7370 |
| 36 | 6R8K-1_A0A0N7KFK3-1 | 0.8004 | 0.7330 |
| 45 | 6R8K-1_Q7XJV3-1 | 0.8290 | 0.7321 |
| 72 | 6R8K-1_Q6YY31-1 | 0.7850 | 0.7303 |
| 140 | 6R8K-1_6R8KA-1* | 0.8007 | 0.7268 |
| 174 | 6R8K-1_Q0JCK8-1 | 0.7562 | 0.7255 |
| 197 | 6R8K-1_Q6EPT2-1 | 0.8069 | 0.7239 |
| 272 | 6R8K-1_Q6EPT4-1 | 0.7668 | 0.7179 |
| 523 | 6R8K-1_D5L9G5-1** | 0.3917 | 0.3387 |

**Supplementary Table 3.** Ranking of complexes of AVR-Pik with true binder and homologous sequences of the true binder in rice. The top scored complexes overall (not homologs) are shaded grey. * Complex with true binder trimmed to only the portion in PDB 6R8K
** Complex with full chain of true binder

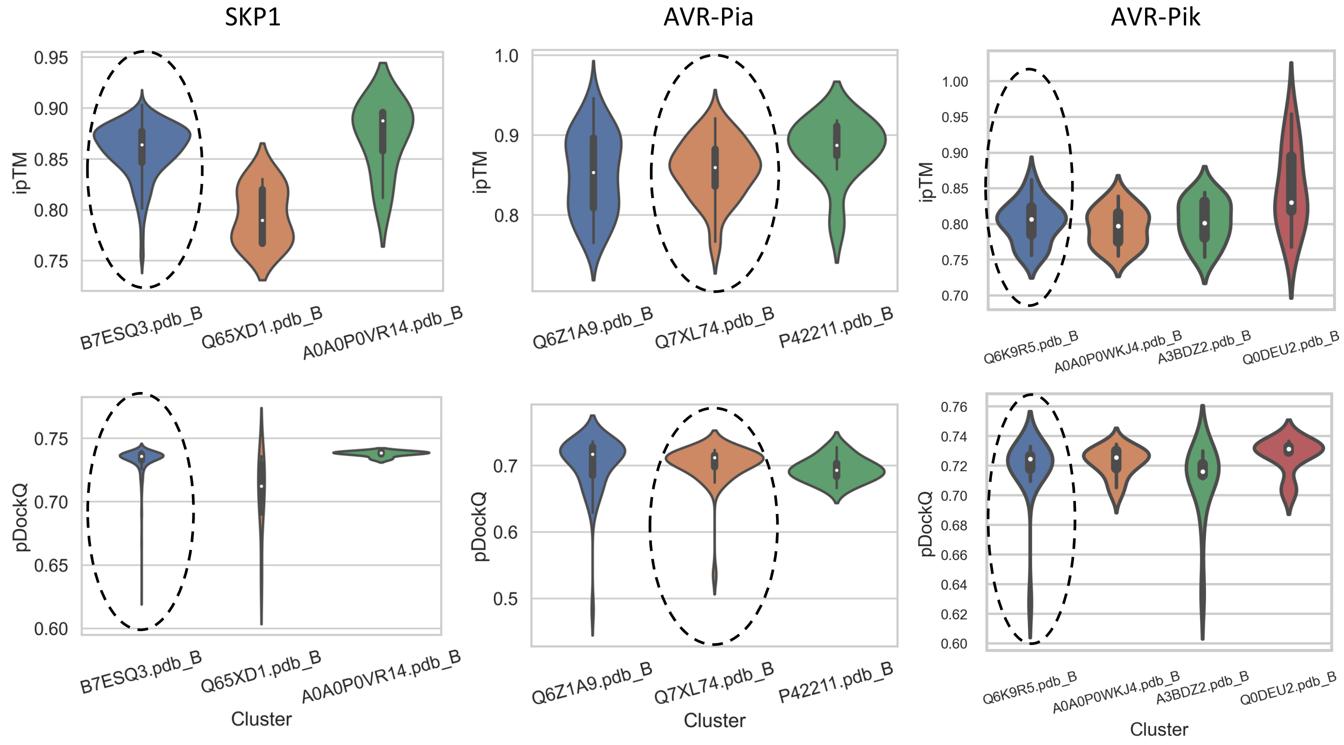


**Supplementary Figure 1.** ipTM and pDockQ scores of the top clusters for the three case studies. The quality scores alone do not provide enough information to identify the clusters with the true binding topologies (circled with dashes).
