## Supplementary Note for "AlphaCRV: A Pipeline for Identifying Accurate Binder Topologies in Mass-Modeling with AlphaFold"

### Clustering SKP1

This notebook will guide you through the process of clustering 712 complexes of the SKP1 protein against a sample of rice proteins. All of the models are dimeric. The 712 models originate from a proteome-wide screen of SKP1 against the rice proteome (O. sativa subsp. japonica, 43,000+ initial models). To select this sample we first ran AlphaCRV on all the models, and the 712 structures are part of some of the best clusters that were identified. These are many more than the other examples because the largest cluster in this case has around 700 structures, which might be due to the higher number of homologues of SKP1 compared to the two AVR proteins in the other examples. Running the pipeline on these models should be enough to reproduce the results and give you an idea of the workflow.

### Prerequisites

- Install AlphaCRV on a conda environment and activate it
- You can download the sequences, resulting clusters, and PyMOL sessions for this example at Zenodo
- The models for this example are very heavy (>140 GB), so we will provide them upon request if you wish to run AlphaCRV on them. Please contact us at, or

### 1. Cluster the models with the `clustering` command

Run the following command to cluster the models. Make sure to change the paths according to your system:

```
alphacrv-cluster \
  --bait ./examples/SKP1/SKP1.fasta \
  --binders ./examples/SKP1/SKP1_binders.fasta \
  --models_dir ./examples/SKP1/SKP1_vs_rice_models \
  --destination ./examples/SKP1/SKP1_vs_rice_clusters \
  --cpus 8
```

After collecting the quality scores from the models in `--models_dir`, it will count how many models are there with an ipTM score higher than the threshold (0.75 by default). It will prompt you to confirm or to modify the threshold. After that, it will proceed with the clustering.

This run will take considerably longer than the two other examples, because of the large size of the first cluster. The full output is as follows:

```
(env) AlphaCRV$ alphacrv-cluster \
--bait ./examples/SKP1/SKP1.fasta \
--binders ./examples/SKP1/SKP1_binders.fasta \
--models_dir ./examples/SKP1/SKP1_vs_rice_models \
--destination ./examples/SKP1/SKP1_vs_rice_clusters \
--cpus 8
INFO:root:Getting quality scores for models in examples/SKP1
/SKP1_vs_rice_models...
INFO:root:Found 712 model directories with quality scores.
Will select 712 models with iptm >= 0.75. Press enter to
continue, or enter a new threshold:
INFO:root:Trimming binder molecules to keep only regions with
an average PAE against the bait of up to 10.0...
INFO:root:Processed 712 complexes.
INFO:root:Writing trimmed sequences to fasta file.
INFO:root:Running sequence clustering...
INFO:root:Processing output...
INFO:root:Running structural clustering...
INFO:root:Processing output...
INFO:root:NOT ALL REPRESENTATIVES COME FROM CHAIN B
INFO:root:Aligning all vs all members of each cluster...
INFO:root:Aligning cluster 1 of 4...
INFO:root:695 members.
INFO:root:Aligning cluster 2 of 4...
INFO:root:6 members.
INFO:root:Aligning cluster 3 of 4...
INFO:root:2 members.
INFO:root:Aligning cluster 4 of 4...
INFO:root:9 members.
INFO:root:Calculating median alignment scores...
INFO:root:Done!!
```

After this step you will have a directory with the following structure:

```
(env) AlphaCRV$ ll examples/SKP1/SKP1_vs_rice_clusters/
total 248K
-rw-r--r-- 1 example g-example  15K Jan  4 13:39
binders_regions.csv
drwxr-xr-x 1 example g-example    0 Jan  4 15:18
merged_clusters/
drwxr-xr-x 1 example g-example    0 Jan  4 13:39
pdbs_trimmed/
drwxr-xr-x 1 example g-example    0 Jan  4 13:39 seqclusters/
drwxr-xr-x 1 example g-example    0 Jan  4 13:39 strclusters/
-rw-r--r-- 1 example g-example 230K Jan  4 13:39
trimmed_binders.fasta
```

Now let's look at some of the important files:

In [ ]:
```
from pathlib import Path
import pandas as pd
```

In [ ]:
```
results_dir = Path('./SKP1/SKP1_vs_rice_clusters/')
```

### See clusters

The `merged_clusters.csv` file contains the list of models with their corresponding sequence, structure and merged clusters. It also has the quality scores provided by AlphaFold.

In [ ]:
```
clusters = pd.read_csv(results_dir / 'merged_clusters/merged_clusters.csv')
```

In [ ]:
```
clusters
```

Out[ ]:

| | complex | str_rep | seq_rep | merged_rep | mer |
|---|---|---|---|---|---|
| **0** | 8IF6-1_Q6Z1A9-1 | A0A0P0Y2U6.pdb_B | A0A0N7KPE6 | A0A0P0XHF4.pdb_B | Q6 |
| **1** | 8IF6-1_Q7XSL8-1 | A0A0P0WQD9.pdb_B | A0A0P0VJA5 | A0A0P0XHF4.pdb_B | Q7 |
| **2** | 8IF6-1_Q5Z8K3-1 | Q6ZDH1.pdb_B | Q5Z8K3 | A0A0P0XHF4.pdb_B | Q5 |
| **3** | 8IF6-1_Q5VMP0-1 | A0A0P0WQD9.pdb_B | Q5VMP0 | A0A0P0XHF4.pdb_B | Q5V |
| **4** | 8IF6-1_A0A0P0WG98-1 | Q5Z7U2.pdb_B | C7J8M0 | A0A0P0XHF4.pdb_B | A0A0P0W |
| **...** | ... | ... | ... | ... | |
| **707** | 8IF6-1_Q0JK63-1 | Q0JK63.pdb_B | Q0JK63 | A0A0P0XHF4.pdb_B | Q0 |
| **708** | 8IF6-1_Q53WL8-1 | Q6ZDH1.pdb_B | A0A0P0WPS2 | A0A0P0XHF4.pdb_B | Q53 |
| **709** | 8IF6-1_A3AQW3-1 | A0A0N7KT42.pdb_B | A3AQW3 | A0A0P0XHF4.pdb_B | A3A |
| **710** | 8IF6-1_Q6ZKB8-1 | Q0DQG8.pdb_B | Q6ZKB8 | A0A0P0XHF4.pdb_B | Q6 |
| **711** | 8IF6-1_Q0DEJ9-1 | A0A0P0XE77.pdb_B | A0A0P0WSK5 | A0A0P0XHF4.pdb_B | Q0 |

712 rows × 7 columns

The columns of the `clusters` DataFrame are:

- `complex` : The name of the complex. This is the same name as the directory where the model is stored.
- `str_rep` : Name of the structure cluster representative
- `seq_rep` : Name of the sequence cluster representative
- `merged_rep` : Name of the merged cluster representative (sequence + structure)
- `member` : The ID of the binder protein
- `iptm` : The ipTM score of the model
- `iptm+ptm` : The ipTM+PTM score of the model (it is calculated by AlphaFold as 0.8*ipTM + 0.2*pTM)

See the number of different merged clusters:

In [ ]:
```
clusters.merged_rep.unique().shape
```

Out[ ]:
```
(4,)
```

For this example, the models and sequences from the 712 binder proteins were summarized in 4 clusters. Much fewer structures to sort through!

### See alignment scores

Alignment scores are calcualted for each cluster by aligning all vs all members of the cluster.

In [ ]:
```
alignment_scores = pd.read_csv(results_dir / 'merged_clusters/alignment_scor
```

In [ ]:
```
alignment_scores.head()
```

Out[ ]:

| | cluster | ref | member | tmscore_ref | tmscore_m | aligned_length | rms |
|---|---|---|---|---|---|---|---|
| 0 | A0A0P0XHF4.pdb_B | Q6Z1A9 | Q7XSL8 | 0.40236 | 0.25976 | 215 | 8.4 |
| 1 | A0A0P0XHF4.pdb_B | Q6Z1A9 | Q5Z8K3 | 0.38146 | 0.28447 | 185 | 10.4 |
| 2 | A0A0P0XHF4.pdb_B | Q6Z1A9 | Q5VMP0 | 0.42881 | 0.23565 | 419 | 16.0 |
| 3 | A0A0P0XHF4.pdb_B | Q6Z1A9 | A0A0P0WG98 | 0.38724 | 0.35182 | 235 | 16.3 |
| 4 | A0A0P0XHF4.pdb_B | Q6Z1A9 | Q67UX0 | 0.40441 | 0.30453 | 263 | 15.2 |

The columns of the `alignment_scores` DataFrame are:

- `cluster` : The name of the cluster
- `ref` : Binder ID of the reference structure in the alignment
- `member` : Binder ID of the second structure in the alignment
- `tmscore_ref` : TM-score based on the reference structure
- `tmscore_m` : TM-score based on the second structure
- `aligned_length` : Length of the alignment
- `rmsd` : RMSD of the alignment

Based on these scores, the median scores are calculated for each cluster member to find the best representative of the cluster (the one with lowest RMSD score to the other members).

### Read median scores and find top clusters

Now we can rank the clusters based on the median alignment scores of the cluster representatives:

In [ ]:
```
median_scores = pd.read_csv(results_dir / 'merged_clusters/median_scores.csv
```

In [ ]:
```
median_scores.shape
```

Out[ ]:
```
(712, 7)
```

The `median_scores` DataFrame contains the median alignment scores of each cluster member when aligned to all other members of the same cluster.

In [ ]:
```
median_scores.head()
```

Out[ ]:

| | cluster | member | tmscore | rmsd | aligned_length | cluster_size | fraction_bi |
|---|---|---|---|---|---|---|---|
| **0** | A0A0P0VR14.pdb_B | A0A0P0VR14 | 0.32871 | 18.64 | 208.0 | 6.0 | 0.11 |
| **1** | A0A0P0VR14.pdb_B | Q2R448 | 0.85991 | 6.04 | 445.0 | 6.0 | 0.93 |
| **2** | A0A0P0VR14.pdb_B | Q6K6K8 | 0.85162 | 4.42 | 445.0 | 6.0 | 0.90 |
| **3** | A0A0P0VR14.pdb_B | Q7XKU0 | 0.49110 | 17.19 | 405.0 | 6.0 | 0.79 |
| **4** | A0A0P0VR14.pdb_B | Q7XL60 | 0.82004 | 4.95 | 455.0 | 6.0 | 0.89 |

The columns of the `median_scores` DataFrame are:

- `cluster` : The name of the cluster
- `member` : ID of the cluster member (binder molecule)
- `tmscore` : Median TM-score of the complex against all other complexes in this cluster
- `rmsd` : Median RMSD of the complex against all other complexes in this cluster
- `aligned_length` : Median length of the alignment
- `cluster_size`
- `fraction_binder` : In average, how much of the binder molecule is included in the alignments of this complex against all other complexes (calculated as `(aligned_length - bait_length)/binder_length` ). This is just meant to be an approximation of how complete the aligmnents are for this cluster member.

The next step is to select the cluster representatives. For this, we first need to filter out the cluster members with poor quality alignments, according to the following criteria:

- Small size
- Low median TM-score
- High median RMSD
- Low fraction of the binder aligned in the cluster representative

In [ ]:

```
# Select the clusters with the following criteria:
select = ((median_scores.cluster_size >= 5) & \
          (median_scores.tmscore >= 0.2) & \
          (median_scores.fraction_binder >= 0.2) & \
          (median_scores.rmsd <= 15))
median_scores_filtered = median_scores[select]
```

In [ ]:

```
median_scores_filtered.shape
```

Out[ ]: (169, 7)

See how many clusters are left after filtering:

In [ ]:

```
median_scores_filtered.cluster.unique().shape
```

Out[ ]: (3,)

Function to format tables:

In [ ]:
```
import seaborn as sns
cm_r = sns.color_palette("mako_r", as_cmap=True)
cm = sns.color_palette("mako", as_cmap=True)
```

In [ ]:
```
def make_pretty(styler):
    styler.format(precision=2)
    styler.background_gradient(axis=0, cmap=cm_r, subset=pd.IndexSlice[:,"cl
    styler.background_gradient(axis=0, cmap=cm_r, subset=pd.IndexSlice[:,"tm
    styler.background_gradient(axis=0, cmap=cm, subset=pd.IndexSlice[:,"rmsd
    styler.background_gradient(axis=0, cmap=cm_r, subset=pd.IndexSlice[:,"fr
    return styler
```

### RESULT 1: See clusters ranked by RMSD

Finally, we can rank the clusters and see which ones have a good combination of low RMSD and large cluster size. These ones are the most likely to contain the true binder.

In [ ]:
```
# Select the rows with the minimum RMSD for each cluster
select = median_scores_filtered.groupby('cluster').rmsd.idxmin()
columns = ['cluster', 'tmscore', 'rmsd', 'cluster_size', 'fraction_binder']
median_scores_filtered.loc[select,columns].sort_values(by='rmsd').style.pipe
```

Out[ ]:

| | cluster | tmscore | rmsd | cluster_size | fraction_binder |
|---|---|---|---|---|---|
| **19** | A0A0P0XHF4.pdb_B | 0.94 | 1.53 | 695.00 | 1.00 |
| **2** | A0A0P0VR14.pdb_B | 0.85 | 4.42 | 6.00 | 0.90 |
| **704** | Q65XD1.pdb_B | 0.70 | 13.05 | 9.00 | 0.93 |

Here we can see that the cluster `B7ESQ3.pdb_B` has the lowest median RMSD and the highest median TM-score. It also has a very large size with 695 members!

### RESULT 2: See clusters ranked by size

In [ ]:
```
# Select the rows with the minimum RMSD for each cluster
select = median_scores_filtered.groupby('cluster').rmsd.idxmin()
columns = ['cluster', 'tmscore', 'rmsd', 'cluster_size', 'fraction_binder']
median_scores_filtered.loc[select, columns].sort_values(by='cluster_size', a
```

Out[ ]:

| | cluster | tmscore | rmsd | cluster_size | fraction_binder |
|---|---|---|---|---|---|
| **19** | A0A0P0XHF4.pdb_B | 0.94 | 1.53 | 695.00 | 1.00 |
| **704** | Q65XD1.pdb_B | 0.70 | 13.05 | 9.00 | 0.93 |
| **2** | A0A0P0VR14.pdb_B | 0.85 | 4.42 | 6.00 | 0.90 |

`B7ESQ3.pdb_B` is also the largest cluster. So we managed to reduce more than 43,000 starting models to only one excellent cluster!

For this example we know that in our list of candidate binders there are 5 homologues of the true binder protein. We can find out which clusters contain these homologues:

In [ ]:
```
homologues = ['Q5VMP0', 'A0A0N7KEW0', 'Q8RZQ3', 'Q69X07', 'A0A0P0Y6A8']
```

In [ ]:
```
clusters[clusters.member.isin(homologues)][['complex','merged_rep','iptm','i
```

Out[ ]:

| | complex | merged_rep | iptm | iptm+ptm |
|---|---|---|---|---|
| **3** | 8IF6-1_Q5VMP0-1 | A0A0P0XHF4.pdb_B | 0.899220 | 0.894956 |
| **28** | 8IF6-1_A0A0P0Y6A8-1 | A0A0P0XHF4.pdb_B | 0.893004 | 0.886242 |
| **321** | 8IF6-1_Q8RZQ3-1 | A0A0P0XHF4.pdb_B | 0.867056 | 0.855604 |
| **372** | 8IF6-1_A0A0N7KEW0-1 | A0A0P0XHF4.pdb_B | 0.862492 | 0.844927 |
| **516** | 8IF6-1_Q69X07-1 | A0A0P0XHF4.pdb_B | 0.847566 | 0.808583 |

They are all in the top cluster!

### 2. Make pymol sessions for the top clusters with `make_pymol_sessions`

Run the following command to select the top clusters that we saw above, make pymol sessions of the top clusters, and optionally do structural clustering on each cluster to find subclusters:

```
alphacrv-rank \
  --clusters_dir ./examples/AVRPia/AVRPia_vs_rice_clusters \
  --min_members 5 \
  --min_tmscore 0.2 \
  --max_rmsd 15 \
  --cluster_clusters
```

The program will show you the top clusters that will be used to make the pymol sessions. You can press `Enter` to continue, or exit the program with `Ctrl+C` to change the filtering parameters.

```
(env) AlphaCRV$ alphacrv-rank \
--clusters_dir ./examples/SKP1/SKP1_vs_rice_clusters \
--min_members 5 \
--min_tmscore 0.2 \
--max_rmsd 15 \
--cluster_clusters
INFO:root:Identified 3 top clusters.
INFO:root:Top clusters:

            cluster   tmscore    rmsd  cluster_size
fraction_binder
0  A0A0P0XHF4.pdb_B  0.941210   1.530         695.0
1.00000
1      Q65XD1.pdb_B  0.704515  13.055           9.0
0.93000
2  A0A0P0VR14.pdb_B  0.851620   4.420           6.0
0.90301

Press Enter to continue, or Ctrl+C to exit and select
different filtering parameters:
INFO:root:Copying pdbs from the top clusters...
INFO:root:Making Pymol sessions...
PyMOL>select chain B AND model A0A0P0VR14_repB
 Selector: selection "sele" defined with 4621 atoms.
PyMOL>bg white
PyMOL>set ray_shadow, 0
 Setting: ray_shadow set to off.
PyMOL>color grey80
 Executive: Colored 42707 atoms.
PyMOL>select chain A
 Selector: selection "sele" defined with 16020 atoms.
PyMOL>color slate, sele
 Executive: Colored 16020 atoms.
PyMOL>delete all
PyMOL>select chain B AND model A0A0P0XHF4_repB
 Selector: selection "sele" defined with 646 atoms.
PyMOL>bg white
PyMOL>set ray_shadow, 0
 Setting: ray_shadow set to off.
PyMOL>color grey80
 Executive: Colored 5270014 atoms.
PyMOL>select chain A
 Selector: selection "sele" defined with 1844949 atoms.
PyMOL>color slate, sele
 Executive: Colored 1844949 atoms.
PyMOL>delete all
PyMOL>select chain B AND model Q65XD1_repB
 Selector: selection "sele" defined with 3154 atoms.
PyMOL>bg white
PyMOL>set ray_shadow, 0
 Setting: ray_shadow set to off.
```

```
PyMOL>color grey80
 Executive: Colored 76932 atoms.
PyMOL>select chain A
 Selector: selection "sele" defined with 24030 atoms.
PyMOL>color slate, sele
 Executive: Colored 24030 atoms.
PyMOL>delete all
INFO:root:Clustering clusters...
INFO:root:Clustering A0A0P0VR14.pdb_B
INFO:root:Running structural clustering...
INFO:root:Processing output...
INFO:root:Clustering A0A0P0XHF4.pdb_B
INFO:root:Running structural clustering...
INFO:root:Processing output...
INFO:root:Clustering Q65XD1.pdb_B
INFO:root:Running structural clustering...
INFO:root:Processing output...
INFO:root:Done!!
```

This command should create the following files in the `./examples/SKP1/SKP1_vs_rice_clusters/merged_clusters/` / `merged_clusters` directory:

- `clustered_clusters.csv` : Contains the subclusters for each of the top clusters.
- `cluster_<cluster_ID>/` : Contains the PDBs of each cluster, and a PyMol session with the cluster members.
- `cluster_<cluster_ID>_clusters/` : Contains the results of the `foldseek easy-cluster` run on the cluster members.

### Read clustered clusters

The following DataFrame contains the subclusters for each of the top clusters:

In [ ]:
```
clustered_clusters = pd.read_csv(results_dir / 'merged_clusters/clustered_cl
```

In [ ]:
```
clustered_clusters.head()
```

Out[ ]:

| | subcluster_rep | member | cluster |
|---|---|---|---|
| 0 | A0A0P0VR14.pdb_B | A0A0P0VR14 | A0A0P0VR14.pdb_B |
| 1 | A0A0P0VR14.pdb_B | Q7XKU0 | A0A0P0VR14.pdb_B |
| 2 | A0A0P0VR14.pdb_B | Q7XL60 | A0A0P0VR14.pdb_B |
| 3 | A0A0P0VR14.pdb_B | Q6K6K8 | A0A0P0VR14.pdb_B |
| 4 | A0A0P0VR14.pdb_B | Q2R448 | A0A0P0VR14.pdb_B |

Now we can look at the most interesting clusters and their subclusters in more detail:

### RESULT 1: Cluster B7ESQ3.pdb_B (contains true binder homologs, top cluster by size)

In [ ]:
```
cluster = 'A0A0P0XHF4.pdb_B'
```

See subclusters:

In [ ]:
```
clustered_clusters[clustered_clusters.cluster==cluster].head()
```

Out[ ]:

| | subcluster_rep | member | cluster |
|---|---|---|---|
| 6 | A0A0P0VGE5.pdb_B | A0A0P0VGE5 | A0A0P0XHF4.pdb_B |
| 7 | A0A0P0VGE5.pdb_B | Q67W96 | A0A0P0XHF4.pdb_B |
| 8 | A0A0P0VHD9.pdb_B | A0A0P0VHD9 | A0A0P0XHF4.pdb_B |
| 9 | A0A0P0VHP5.pdb_B | A0A0P0VHP5 | A0A0P0XHF4.pdb_B |
| 10 | A0A0P0VHP5.pdb_B | A0A0P0UZC1 | A0A0P0XHF4.pdb_B |

Count how many subclusters:

In [ ]:
```
clustered_clusters[clustered_clusters.cluster==cluster].subcluster_rep.uniqu
```

Out[ ]:
```
(53,)
```

See the amount of strucures in each subcluster:

In [ ]:
```
(clustered_clusters[clustered_clusters.cluster==cluster].groupby('subcluster
 .size().sort_values(ascending=False).head(15))
```

Out[ ]:
```
subcluster_rep
Q6ZDH1.pdb_B        223
A0A0P0WQD9.pdb_B    116
A0A0P0Y2U6.pdb_B     51
Q5Z7U2.pdb_B         35
Q0DQG8.pdb_B         34
Q75J50.pdb_B         27
Q0JPJ6.pdb_B         19
A0A0P0XHI5.pdb_B     14
A0A0P0X3V6.pdb_B     14
A0A0P0VHP5.pdb_B     12
Q7XAK4.pdb_B          9
Q7EY32.pdb_B          9
A0A0P0V3R7.pdb_B      9
Q8LJA9.pdb_B          8
A0A0P0X458.pdb_B      7
dtype: int64
```

Here we have 53 subclusters!! This cluster has in total 695 strucures, which is a bit too much to load into PyMOL at once. We have to change our strategy. One way to decide if the subclusters are similar or different to each other would be to only look at the subcluster representatives and classify them according to their topology. We will try now three different approaches to visualize this cluster:

### Take all members of this cluster together

We can just look at the structures with lowest median RMSD to have an idea of how this cluster looks:

In [ ]:
```
median_scores[median_scores.cluster==cluster].sort_values(by='rmsd').head()
```

Out[ ]:

| | cluster | member | tmscore | rmsd | aligned_length | cluster_size | fractio |
|---|---|---|---|---|---|---|---|
| **19** | A0A0P0XHF4.pdb_B | A0A0N7KK85 | 0.941210 | 1.530 | 210.0 | 695.0 | |
| **520** | A0A0P0XHF4.pdb_B | Q6ASY4 | 0.938210 | 1.560 | 213.0 | 695.0 | |
| **401** | A0A0P0XHF4.pdb_B | Q2QQH8 | 0.938700 | 1.600 | 215.0 | 695.0 | |
| **37** | A0A0P0XHF4.pdb_B | A0A0N7KT42 | 0.935435 | 1.630 | 208.0 | 695.0 | |
| **256** | A0A0P0XHF4.pdb_B | A0A0P0YAQ4 | 0.935185 | 1.645 | 217.0 | 695.0 | |

We can print out the names of the top 20 models to open them in PyMOL (through the terminal or a command inside of PyMOL)

In [ ]:
```
path_cluster = Path.cwd().resolve()
```

In [ ]:
```
path_cluster = '/Volumes/weka_user/guzmanfj/py/AlphaCRV/examples/'
```

In [ ]:
```
# Print paths to open models in pymol
membs = median_scores[median_scores.cluster==cluster].sort_values(by='rmsd')
' '.join([(path_cluster + str(results_dir) + f'/merged_clusters/cluster_{clu
```

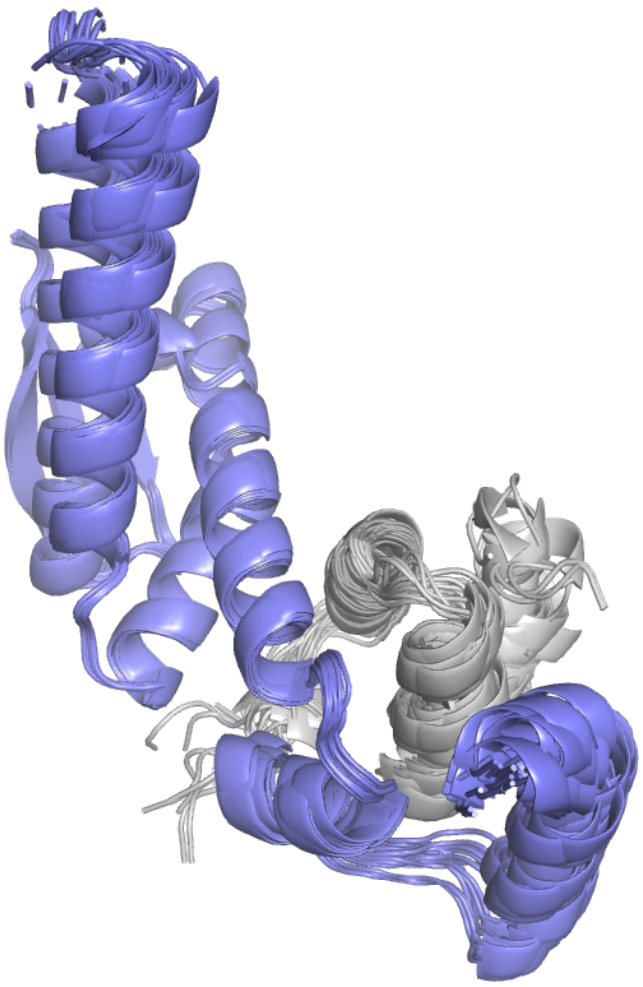

### Look at all the subcluster representatives in PyMOL and group them by topology.

In [ ]:
```
subclusters = (clustered_clusters[clustered_clusters.cluster==cluster])
subclusters_ids = subclusters['subcluster_rep'].str.split('.pdb').str[0]
```

See the names of the subclusters:

In [ ]:
```
subclusters.subcluster_rep.unique()
```

```
Out[ ]: array(['A0A0P0VGE5.pdb_B', 'A0A0P0VHD9.pdb_B', 'A0A0P0VHP5.pdb_B',
               'A0A0P0VNF2.pdb_B', 'A0A0P0VNI4.pdb_B', 'A0A0P0VRN2.pdb_B',
               'A0A0P0VZH9.pdb_B', 'A0A0P0W419.pdb_B', 'A0A0P0X4D4.pdb_B',
               'A0A0P0XCY3.pdb_B', 'A0A0P0XE00.pdb_B', 'A0A0P0XE77.pdb_B',
               'A0A0P0XEW7.pdb_B', 'A0A0P0XFT5.pdb_B', 'A0A0P0XHF4.pdb_B',
               'A0A0P0XHI5.pdb_B', 'A0A0P0XNQ3.pdb_B', 'A0A0P0XSL8.pdb_B',
               'A0A0P0Y2U6.pdb_B', 'A0A0P0Y5S9.pdb_B', 'A0A0P0YA59.pdb_B',
               'A0A0P0YAA2.pdb_B', 'A0A0P0YC15.pdb_B', 'A2ZPC4.pdb_B',
               'B7ESQ3.pdb_B', 'B9G6M6.pdb_B', 'Q0DQG8.pdb_B', 'Q0JB10.pdb_B',
               'Q0JK63.pdb_B', 'Q0JPJ6.pdb_B', 'Q109D8.pdb_B', 'Q2QQH8.pdb_B',
               'Q6ZDH1.pdb_B', 'A0A0N7KD81.pdb_A', 'A0A0N7KK85.pdb_B',
               'A0A0N7KN01.pdb_B', 'A0A0N7KT42.pdb_B', 'A0A0P0V091.pdb_B',
               'A0A0P0V3R7.pdb_B', 'A0A0P0WDY8.pdb_B', 'A0A0P0WQD9.pdb_B',
               'A0A0P0X3V6.pdb_B', 'A0A0P0X458.pdb_B', 'Q5N762.pdb_B',
               'Q5Z7U2.pdb_B', 'Q69J07.pdb_B', 'Q69X07.pdb_B', 'Q75IQ5.pdb_B',
               'Q75J50.pdb_B', 'Q7EY32.pdb_B', 'Q7FAH1.pdb_B', 'Q7XAK4.pdb_B',
               'Q8LJA9.pdb_B'], dtype=object)
```

Print the PATHs of the subcluster representatives to open them in PyMOL:

```
In [ ]: ' '.join([path_cluster + str(results_dir) + f'/merged_clusters/cluster_{clus
```

Write down the subclusters in different categories:

Binding motif --- A0A0P0VGE5 A0A0P0VHD9 A0A0P0VHP5 A0A0P0VNF2 A0A0P0VNI4 A0A0P0X4D4 A0A0P0XCY3 A0A0P0XE00 A0A0P0XE77 A0A0P0XEW7 A0A0P0XHF4 A0A0P0XHI5 A0A0P0XNQ3 A0A0P0XSL8 A0A0P0YA59 A0A0P0YAA2 B7ESQ3 B9G6M6 Q109D8 Q2QQH8 A0A0N7KK85 A0A0N7KT42 A0A0P0X458 Q5N762 Q69J07 Q69X07 Q75IQ5 Q7EY32 Q7XAK4 Horseshoe --- A0A0P0VRN2 A0A0P0XFT5 A0A0P0Y2U6 A0A0P0Y5S9 A0A0P0YC15 A0A0N7KD81 A0A0P0V091 A0A0P0WQD9 A0A0P0X3V6 Q5Z7U2 Q75J50 Q7FAH1 Beta propeller --- A0A0P0W419 Q0DQG8 Q0JK63 Q6ZDH1 A0A0P0V3R7 Beta other --- A0A0P0VZH9 A2ZPC4 Q0JB10 Q0JPJ6 A0A0N7KN01 A0A0P0WDY8 Q8LJA9

### Subclusters for only the binding motif

In [ ]:
```
subcluster_names = [
'A0A0P0VGE5.pdb_B',
'A0A0P0VHD9.pdb_B',
'A0A0P0VHP5.pdb_B',
'A0A0P0VNF2.pdb_B',
'A0A0P0VNI4.pdb_B',
'A0A0P0X4D4.pdb_B',
'A0A0P0XCY3.pdb_B',
'A0A0P0XE00.pdb_B',
'A0A0P0XE77.pdb_B',
'A0A0P0XEW7.pdb_B',
'A0A0P0XHF4.pdb_B',
'A0A0P0XHI5.pdb_B',
'A0A0P0XNQ3.pdb_B',
'A0A0P0XSL8.pdb_B',
'A0A0P0YA59.pdb_B',
'A0A0P0YAA2.pdb_B',
'B7ESQ3.pdb_B',
'B9G6M6.pdb_B',
'Q109D8.pdb_B',
'Q2QQH8.pdb_B',
'A0A0N7KK85.pdb_B',
'A0A0N7KT42.pdb_B',
'A0A0P0X458.pdb_B',
'Q5N762.pdb_B',
'Q69J07.pdb_B',
'Q69X07.pdb_B',
'Q75IQ5.pdb_B',
'Q7EY32.pdb_B',
'Q7XAK4.pdb_B'
]
```

See how many proteins are in these subclusters:

In [ ]:
```
(clustered_clusters[clustered_clusters.subcluster_rep.isin(subcluster_names)
 .subcluster_rep.value_counts().sum())
```

Out[ ]: 113

See the alignment scores of the top members in these subclusters:

In [ ]:
```
# Get list of members for subcluster
members_sc = list(clustered_clusters[clustered_clusters.subcluster_rep.isin(
                  .member)

median_scores[median_scores.member.isin(members_sc)].sort_values(by='rmsd').
```

Out[ ]:

| | cluster | member | tmscore | rmsd | aligned_length | cluster_size | fracti |
|---|---|---|---|---|---|---|---|
| **19** | A0A0P0XHF4.pdb_B | A0A0N7KK85 | 0.941210 | 1.530 | 210.0 | 695.0 | |
| **520** | A0A0P0XHF4.pdb_B | Q6ASY4 | 0.938210 | 1.560 | 213.0 | 695.0 | |
| **401** | A0A0P0XHF4.pdb_B | Q2QQH8 | 0.938700 | 1.600 | 215.0 | 695.0 | |
| **37** | A0A0P0XHF4.pdb_B | A0A0N7KT42 | 0.935435 | 1.630 | 208.0 | 695.0 | |
| **256** | A0A0P0XHF4.pdb_B | A0A0P0YAQ4 | 0.935185 | 1.645 | 217.0 | 695.0 | |
| **700** | A0A0P0XHF4.pdb_B | Q9FWD4 | 0.923980 | 1.740 | 216.0 | 695.0 | |
| **418** | A0A0P0XHF4.pdb_B | Q2R0S7 | 0.934600 | 1.760 | 217.0 | 695.0 | |
| **145** | A0A0P0XHF4.pdb_B | A0A0P0X0M4 | 0.921310 | 1.780 | 215.0 | 695.0 | |
| **290** | A0A0P0XHF4.pdb_B | B9FCI9 | 0.919485 | 1.830 | 219.0 | 695.0 | |
| **394** | A0A0P0XHF4.pdb_B | Q2QMT0 | 0.918475 | 1.830 | 216.0 | 695.0 | |
| **88** | A0A0P0XHF4.pdb_B | A0A0P0W0W5 | 0.933120 | 1.840 | 218.0 | 695.0 | |
| **14** | A0A0P0XHF4.pdb_B | A0A0N7KHR2 | 0.932530 | 1.850 | 218.0 | 695.0 | |
| **480** | A0A0P0XHF4.pdb_B | Q5VNR1 | 0.922670 | 1.850 | 216.0 | 695.0 | |
| **333** | A0A0P0XHF4.pdb_B | Q0ITY1 | 0.917245 | 1.855 | 206.0 | 695.0 | |
| **693** | A0A0P0XHF4.pdb_B | Q943F5 | 0.924290 | 1.860 | 213.0 | 695.0 | |
| **361** | A0A0P0XHF4.pdb_B | Q0JI48 | 0.924270 | 1.870 | 218.0 | 695.0 | |
| **71** | A0A0P0XHF4.pdb_B | A0A0P0VLY9 | 0.923905 | 1.880 | 217.0 | 695.0 | |
| **232** | A0A0P0XHF4.pdb_B | A0A0P0XZS4 | 0.921440 | 1.910 | 218.0 | 695.0 | |
| **157** | A0A0P0XHF4.pdb_B | A0A0P0X4D4 | 0.915655 | 1.915 | 212.0 | 695.0 | |
| **254** | A0A0P0XHF4.pdb_B | A0A0P0YAA2 | 0.914570 | 1.940 | 211.0 | 695.0 | |

See if any of the homologous proteins are in here:

In [ ]:
```
[h in members_sc for h in homologues]
```

Out[ ]:
```
[False, False, False, True, False]
```

In [ ]:
```
homologues
```

Out[ ]:
```
['Q5VMP0', 'A0A0N7KEW0', 'Q8RZQ3', 'Q69X07', 'A0A0P0Y6A8']
```

`Q69X07` is in this subcluster.

Print the paths of the top 20 PDB files to open them in PyMOL:

In [ ]:
```
# Print paths to open models in pymol
membs = median_scores[median_scores.member.isin(members_sc)].sort_values(by=
' '.join([(path_cluster + str(results_dir) + f'/merged_clusters/cluster_{clu
```

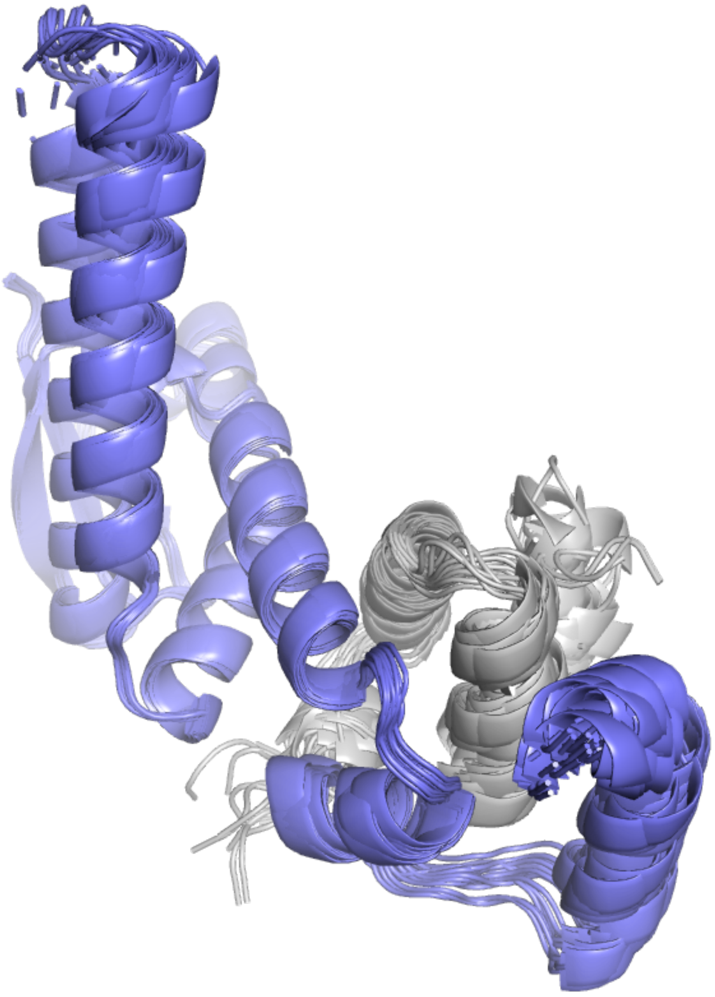

This is very similar to the previous figure with all the subclusters together. The models with the lowest RMSD overall are those who only contain the minimal bindign motif of the binder protein, since they have fewer residues to align.

#### Subclusters for the binding motif + Leucine-rich region (horseshoe)

In [ ]:
```
subcluster_names = [
'A0A0P0VRN2.pdb_B',
'A0A0P0XFT5.pdb_B',
'A0A0P0Y2U6.pdb_B',
'A0A0P0Y5S9.pdb_B',
'A0A0P0YC15.pdb_B',
'A0A0N7KD81.pdb_B',
'A0A0P0V091.pdb_B',
'A0A0P0WQD9.pdb_B',
'A0A0P0X3V6.pdb_B',
'Q5Z7U2.pdb_B',
'Q75J50.pdb_B',
'Q7FAH1.pdb_B'
]
```

See how many proteins are in these subclusters:

In [ ]:
```
(clustered_clusters[clustered_clusters.subcluster_rep.isin(subcluster_names)
 .subcluster_rep.value_counts().sum())
```

Out[ ]: 262

See the alignment scores of the top members in these subclusters:

In [ ]:
```
# Get list of members for subcluster
members_sc = list(clustered_clusters[clustered_clusters.subcluster_rep.isin(
                  .member)

median_scores[median_scores.member.isin(members_sc)].sort_values(by='rmsd').
```

Out[ ]:

| | cluster | member | tmscore | rmsd | aligned_length | cluster_size | fracti |
|---|---|---|---|---|---|---|---|
| **171** | A0A0P0XHF4.pdb_B | A0A0P0XBZ2 | 0.687465 | 7.325 | 230.5 | 695.0 | |
| **149** | A0A0P0XHF4.pdb_B | A0A0P0X2K4 | 0.415300 | 7.340 | 216.0 | 695.0 | |
| **82** | A0A0P0XHF4.pdb_B | A0A0P0VUW0 | 0.711865 | 7.670 | 270.0 | 695.0 | |
| **413** | A0A0P0XHF4.pdb_B | Q2QY97 | 0.432800 | 7.685 | 221.0 | 695.0 | |
| **81** | A0A0P0XHF4.pdb_B | A0A0P0VRP6 | 0.596455 | 7.750 | 221.5 | 695.0 | |
| **310** | A0A0P0XHF4.pdb_B | C7J8M0 | 0.600700 | 7.785 | 219.0 | 695.0 | |
| **118** | A0A0P0XHF4.pdb_B | A0A0P0WF72 | 0.641515 | 7.825 | 238.0 | 695.0 | |
| **282** | A0A0P0XHF4.pdb_B | B7EXZ6 | 0.514025 | 8.010 | 214.0 | 695.0 | |
| **249** | A0A0P0XHF4.pdb_B | A0A0P0Y5S9 | 0.551080 | 8.015 | 209.0 | 695.0 | |
| **244** | A0A0P0XHF4.pdb_B | A0A0P0Y2Y9 | 0.417340 | 8.035 | 218.5 | 695.0 | |
| **367** | A0A0P0XHF4.pdb_B | Q0JLI9 | 0.285820 | 8.055 | 223.0 | 695.0 | |
| **359** | A0A0P0XHF4.pdb_B | Q0JES5 | 0.269000 | 8.165 | 217.5 | 695.0 | |
| **366** | A0A0P0XHF4.pdb_B | Q0JLI0 | 0.276000 | 8.175 | 220.0 | 695.0 | |
| **69** | A0A0P0XHF4.pdb_B | A0A0P0VJA5 | 0.375275 | 8.200 | 230.0 | 695.0 | |
| **248** | A0A0P0XHF4.pdb_B | A0A0P0Y4Y4 | 0.704520 | 8.275 | 242.5 | 695.0 | |
| **647** | A0A0P0XHF4.pdb_B | Q7XSL8 | 0.264890 | 8.320 | 222.0 | 695.0 | |
| **34** | A0A0P0XHF4.pdb_B | A0A0N7KR34 | 0.508930 | 8.380 | 215.0 | 695.0 | |
| **358** | A0A0P0XHF4.pdb_B | Q0JES4 | 0.275240 | 8.555 | 218.5 | 695.0 | |
| **533** | A0A0P0XHF4.pdb_B | Q6ETW1 | 0.453605 | 8.555 | 222.5 | 695.0 | |
| **667** | A0A0P0XHF4.pdb_B | Q8GVM0 | 0.290510 | 8.570 | 220.5 | 695.0 | |

See if any of the homologous proteins are in here:

In [ ]:

```
[h in members_sc for h in homologues]
```

Out[ ]:

```
[True, True, True, False, True]
```

In [ ]:

```
homologues
```

Out[ ]:

```
['Q5VMP0', 'A0A0N7KEW0', 'Q8RZQ3', 'Q69X07', 'A0A0P0Y6A8']
```

All the rest of homologous proteins are in here!

Print the paths of the top 20 PDB files to open them in PyMOL:

In [ ]:

```
# Print paths to open models in pymol
membs = median_scores[median_scores.member.isin(members_sc)].sort_values(by=
' '.join([(path_cluster + str(results_dir) + f'/merged_clusters/cluster_{clu
```

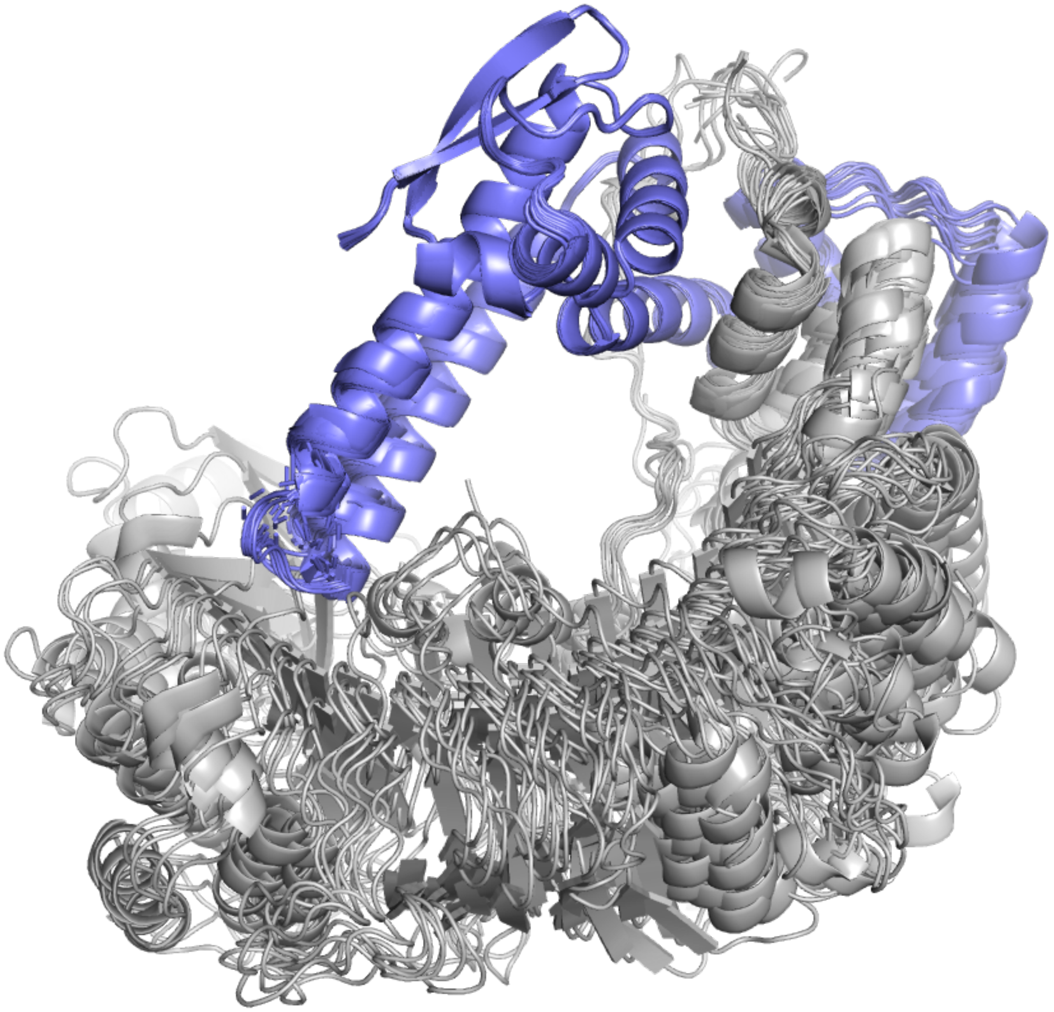

These subclusters contain binders that, in addition to the binding motif, have LLR domains that also interact with SKP1 in some way. The true binder of SKP1 and its homologues have a similar architecture.

### Subclusters for the binding motif + beta propeller

In [ ]:
```
subcluster_names = [
'A0A0P0W419.pdb_B',
'Q0DQG8.pdb_B',
'Q0JK63.pdb_B',
'Q6ZDH1.pdb_B',
'A0A0P0V3R7.pdb_B'
]
```

See how many proteins are in these subclusters:

In [ ]:
```
(clustered_clusters[clustered_clusters.subcluster_rep.isin(subcluster_names)
 .subcluster_rep.value_counts().sum())
```

Out[ ]: 277

See the alignment scores of the top members in these subclusters:

In [ ]:
```
# Get list of members for subcluster
members_sc = list(clustered_clusters[clustered_clusters.subcluster_rep.isin(
                  .member)

median_scores[median_scores.member.isin(members_sc)].sort_values(by='rmsd').
```

Out[ ]:

| | cluster | member | tmscore | rmsd | aligned_length | cluster_size | fracti |
|---|---|---|---|---|---|---|---|
| **213** | A0A0P0XHF4.pdb_B | A0A0P0XQK8 | 0.302510 | 7.940 | 201.5 | 695.0 | |
| **133** | A0A0P0XHF4.pdb_B | A0A0P0WR08 | 0.309095 | 8.335 | 188.0 | 695.0 | |
| **422** | A0A0P0XHF4.pdb_B | Q2R1S8 | 0.293405 | 8.740 | 194.0 | 695.0 | |
| **528** | A0A0P0XHF4.pdb_B | Q6EQC5 | 0.351750 | 8.760 | 197.5 | 695.0 | |
| **575** | A0A0P0XHF4.pdb_B | Q6Z4S1 | 0.318150 | 8.760 | 206.5 | 695.0 | |
| **691** | A0A0P0XHF4.pdb_B | Q8VWI8 | 0.295920 | 8.865 | 194.0 | 695.0 | |
| **360** | A0A0P0XHF4.pdb_B | Q0JFD5 | 0.274020 | 8.890 | 191.0 | 695.0 | |
| **692** | A0A0P0XHF4.pdb_B | Q8W0I3 | 0.308530 | 8.955 | 195.0 | 695.0 | |
| **471** | A0A0P0XHF4.pdb_B | Q53WL8 | 0.333390 | 9.060 | 197.0 | 695.0 | |
| **577** | A0A0P0XHF4.pdb_B | Q6Z6Y9 | 0.307905 | 9.070 | 197.0 | 695.0 | |
| **66** | A0A0P0XHF4.pdb_B | A0A0P0VI66 | 0.302140 | 9.100 | 194.0 | 695.0 | |
| **129** | A0A0P0XHF4.pdb_B | A0A0P0WQ78 | 0.293750 | 9.125 | 191.0 | 695.0 | |
| **599** | A0A0P0XHF4.pdb_B | Q6ZK55 | 0.313985 | 9.145 | 198.0 | 695.0 | |
| **578** | A0A0P0XHF4.pdb_B | Q6Z7A7 | 0.333375 | 9.185 | 197.0 | 695.0 | |
| **513** | A0A0P0XHF4.pdb_B | Q69RN5 | 0.279535 | 9.300 | 192.0 | 695.0 | |
| **474** | A0A0P0XHF4.pdb_B | Q5QM20 | 0.326920 | 9.310 | 213.0 | 695.0 | |
| **128** | A0A0P0XHF4.pdb_B | A0A0P0WPS2 | 0.290210 | 9.330 | 195.0 | 695.0 | |
| **267** | A0A0P0XHF4.pdb_B | A0A5S6R9A0 | 0.329460 | 9.330 | 209.0 | 695.0 | |
| **677** | A0A0P0XHF4.pdb_B | Q8LHX9 | 0.326415 | 9.380 | 202.5 | 695.0 | |
| **363** | A0A0P0XHF4.pdb_B | Q0JKP0 | 0.297210 | 9.410 | 195.0 | 695.0 | |

Print the paths of the top 20 PDB files to open them in PyMOL:

In [ ]:
```
# Print paths to open models in pymol
membs = median_scores[median_scores.member.isin(members_sc)].sort_values(by=
' '.join([(path_cluster + str(results_dir) + f'/merged_clusters/cluster_{clu
```

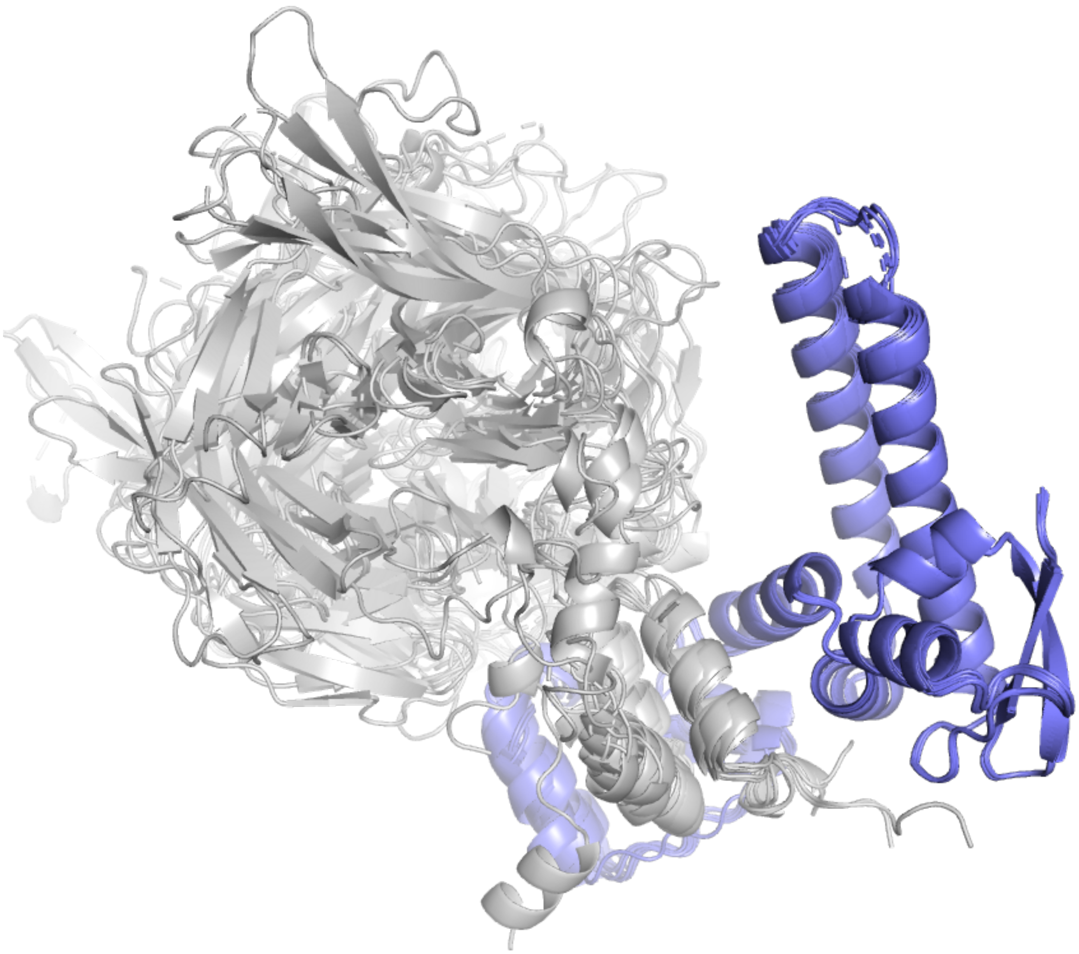

The alignment of the beta propeller domains is much messier, as they are modeled in many different orientations. The screenshot above only contains a few of the complexes.

In [ ]:

### Clustering AVR-Pia

This notebook will guide you through the process of clustering 99 complexes of the AVR-Pia protein against a sample of rice proteins. All of the models are dimeric. The 99 models originate from a proteome-wide screen of AVR-Pia against the rice proteome (O. sativa subsp. japonica, 43,000+ initial models). To select this sample we first ran AlphaCRV on all the models, and the 99 structures are part of some of the best clusters that were identified. Running the pipeline on these models should be enough to reproduce the results and give you an idea of the workflow.

### Prerequisites

- Install AlphaCRV on a conda environment and activate it
- Download the sequences, models and results for this example at Zenodo

### 1. Cluster the models with the `clustering` command

Run the following command to cluster the models. Make sure to change the paths according to your system:

```
alphacrv-cluster \
  --bait ./examples/AVRPia/AVRPia.fasta \
  --binders ./examples/AVRPia/AVRPia_binders.fasta \
  --models_dir ./examples/AVRPia/AVRPia_vs_rice_models \
  --destination ./examples/AVRPia/AVRPia_vs_rice_clusters \
  --cpus 8
```

After collecting the quality scores from the models in `--models_dir` , it will count how many models are there with an ipTM scores higher than the threshold (0.75 by default). It will prompt you to confirm or to modify the threshold. After that, it will proceed with the clustering.

The full output is as follows:

```
(env) AlphaCRV$ alphacrv-cluster \
--bait ./examples/AVRPia/AVRPia.fasta \
--binders ./examples/AVRPia/AVRPia_binders.fasta \
--models_dir ./examples/AVRPia/AVRPia_vs_rice_models \
--destination ./examples/AVRPia/AVRPia_vs_rice_clusters \
--cpus 8
INFO:root:Getting quality scores for models in
examples/AVRPia/AVRPia_vs_rice_models...
INFO:root:Found 99 model directories with quality scores.
Will select 99 models with iptm >= 0.75. Press enter to
continue, or enter a new threshold:
INFO:root:Trimming binder molecules to keep only regions with
an average PAE against the bait of up to 10.0...
INFO:root:Processed 99 complexes.
INFO:root:Writing trimmed sequences to fasta file.
INFO:root:Running sequence clustering...
INFO:root:Processing output...
INFO:root:Running structural clustering...
INFO:root:Processing output...
INFO:root:Aligning all vs all members of each cluster...
INFO:root:Aligning cluster 1 of 3...
INFO:root:49 members.
INFO:root:Aligning cluster 2 of 3...
INFO:root:40 members.
INFO:root:Aligning cluster 3 of 3...
INFO:root:10 members.
INFO:root:Calculating median alignment scores...
INFO:root:Done!!
```

After this step you will have a directory with the following structure:

```
(env) AlphaCRV$ ll examples/AVRPia/AVRPia_vs_rice_clusters/
total 36K
-rw-r--r-- 1 example g-example 2.0K Jan  3 19:39
binders_regions.csv
drwxr-xr-x 1 example g-example    0 Jan  3 19:40
merged_clusters/
drwxr-xr-x 1 example g-example    0 Jan  3 19:39
pdbs_trimmed/
drwxr-xr-x 1 example g-example    0 Jan  3 19:39 seqclusters/
drwxr-xr-x 1 example g-example    0 Jan  3 19:39 strclusters/
-rw-r--r-- 1 example g-example  29K Jan  3 19:39
trimmed_binders.fasta
```

Now let's look at some of the important files:

In [ ]:
```
from pathlib import Path
import pandas as pd
```

In [ ]:
```
results_dir = Path('./AVRPia/AVRPia_vs_rice_clusters/')
```

#### See clusters

The `merged_clusters.csv` file contains the list of models with their corresponding sequence, structure and merged clusters. It also has the quality scores provided by AlphaFold.

In [ ]:
```
clusters = pd.read_csv(results_dir / 'merged_clusters/merged_clusters.csv')
```

In [ ]:
```
clusters
```

Out[ ]:

| | complex | str_rep | seq_rep | merged_rep | member |
|---|---|---|---|---|---|
| **0** | 6Q76-1_Q5JN27-1 | Q0IYV8.pdb_B | C7IWS6 | Q6Z1A9.pdb_B | Q5JN27 |
| **1** | 6Q76-1_Q6K9S6-1 | A0A0P0WCR3.pdb_B | Q6K9S6 | Q6Z1A9.pdb_B | Q6K9S6 |
| **2** | 6Q76-1_A0A0P0VEX0-1 | Q0IYV8.pdb_B | A0A0P0XZP1 | Q6Z1A9.pdb_B | A0A0P0VEX0 |
| **3** | 6Q76-1_Q2R3L3-1 | A0A0P0VF33.pdb_B | A0A0P0Y2U8 | Q6Z1A9.pdb_B | Q2R3L3 |
| **4** | 6Q76-1_A0A0P0UZA6-1 | A0A0P0UZA6.pdb_B | A0A0P0UZA6 | Q7XL74.pdb_B | A0A0P0UZA6 |
| **...** | ... | ... | ... | ... | ... |
| **94** | 6Q76-1_A0A0P0XDF8-1 | Q0IYV8.pdb_B | A0A0N7KMC1 | Q6Z1A9.pdb_B | A0A0P0XDF8 |
| **95** | 6Q76-1_A0A0P0Y3K7-1 | Q0IYV8.pdb_B | A0A0P0Y3K7 | Q6Z1A9.pdb_B | A0A0P0Y3K7 |
| **96** | 6Q76-1_Q0IYI4-1 | Q7XL74.pdb_B | Q7XL74 | Q7XL74.pdb_B | Q0IYI4 |
| **97** | 6Q76-1_Q6Z0A9-1 | Q0IYV8.pdb_B | A0A0N7KMC1 | Q6Z1A9.pdb_B | Q6Z0A9 |
| **98** | 6Q76-1_Q84TS2-1 | Q7XL74.pdb_B | Q8S181 | Q7XL74.pdb_B | Q84TS2 |

99 rows × 7 columns

The columns of the `clusters` DataFrame are:

- `complex` : The name of the complex. This is the same name as the directory where the model is stored.
- `str_rep` : Name of the structure cluster representative
- `seq_rep` : Name of the sequence cluster representative
- `merged_rep` : Name of the merged cluster representative (sequence + structure)
- `member` : The ID of the binder protein
- `iptm` : The ipTM score of the model
- `iptm+ptm` : The ipTM+PTM score of the model (it is calculated by AlphaFold as 0.8*ipTM + 0.2*pTM)

See the number of different merged clusters:

In [ ]:
```
clusters.merged_rep.unique().shape
```

Out[ ]: (3,)

For this example, the models and sequences from the 99 binder proteins were summarized in 3 clusters. Much fewer structures to sort through!

### See alignment scores

Alignment scores are calcualted for each cluster by aligning all vs all members of the cluster.

In [ ]:
```
alignment_scores = pd.read_csv(results_dir / 'merged_clusters/alignment_scor
```

In [ ]:
```
alignment_scores.head()
```

Out[ ]:

| | cluster | ref | member | tmscore_ref | tmscore_m | aligned_length | rmsd |
|---|---|---|---|---|---|---|---|
| **0** | Q6Z1A9.pdb_B | Q5JN27 | Q6K9S6 | 0.22351 | 0.56157 | 206 | 12.72 |
| **1** | Q6Z1A9.pdb_B | Q5JN27 | A0A0P0VEX0 | 0.53580 | 0.44881 | 564 | 20.63 |
| **2** | Q6Z1A9.pdb_B | Q5JN27 | Q2R3L3 | 0.32890 | 0.66186 | 274 | 10.16 |
| **3** | Q6Z1A9.pdb_B | Q5JN27 | A0A0P0Y2U8 | 0.23944 | 0.51955 | 225 | 11.05 |
| **4** | Q6Z1A9.pdb_B | Q5JN27 | C7IWS6 | 0.55887 | 0.81560 | 393 | 9.61 |

The columns of the `alignment_scores` DataFrame are:

- `cluster` : The name of the cluster
- `ref` : Binder ID of the reference structure in the alignment
- `member` : Binder ID of the second structure in the alignment
- `tmscore_ref` : TM-score based on the reference structure
- `tmscore_m` : TM-score based on the second structure
- `aligned_length` : Length of the alignment
- `rmsd` : RMSD of the alignment

Based on these scores, the median scores are calculated for each cluster member to find the best representative of the cluster (the one with lowest RMSD score to the other members).

### Read median scores and find top clusters

Now we can rank the clusters based on the median alignment scores of the cluster representatives:

In [ ]:
```
median_scores = pd.read_csv(results_dir / 'merged_clusters/median_scores.csv
```

In [ ]:
```
median_scores.shape
```

Out[ ]: (99, 7)

The `median_scores` DataFrame contains the median alignment scores of each cluster member when aligned to all other members of the same cluster.

In [ ]:
```
median_scores.head()
```

Out[ ]:

| | cluster | member | tmscore | rmsd | aligned_length | cluster_size | fraction_binder |
|---|---|---|---|---|---|---|---|
| **0** | P42211.pdb_B | A0A0P0VAB8 | 0.56928 | 21.55 | 419.0 | 10.0 | 0.862408 |
| **1** | P42211.pdb_B | A0A0P0VAW6 | 0.59934 | 12.81 | 407.0 | 10.0 | 0.867008 |
| **2** | P42211.pdb_B | A0A0P0WVN9 | 0.39349 | 24.73 | 412.0 | 10.0 | 0.577181 |
| **3** | P42211.pdb_B | P42211 | 0.26818 | 20.91 | 363.0 | 10.0 | 0.707434 |
| **4** | P42211.pdb_B | Q0D5V1 | 0.66389 | 10.75 | 402.0 | 10.0 | 0.902703 |

The columns of the `median_scores` DataFrame are:

- `cluster` : The name of the cluster
- `member` : ID of the cluster member (binder molecule)
- `tmscore` : Median TM-score of the complex against all other complexes in this cluster
- `rmsd` : Median RMSD of the complex against all other complexes in this cluster
- `aligned_length` : Median length of the alignment
- `cluster_size`
- `fraction_binder` : In average, how much of the binder molecule is included in the alignments of this complex against all other complexes (calculated as `(aligned_length - bait_length)/binder_length` ). This is just meant to be an approximation of how complete the aligmnents are for this cluster member.

The next step is to select the cluster representatives. For this, we first need to filter out the cluster members with poor quality alignments, according to the following criteria:

- Small size
- Low median TM-score
- High median RMSD
- Low fraction of the binder aligned in the cluster representative

In [ ]:
```
# Select the clusters with the following criteria:
select = ((median_scores.cluster_size >= 5) & \
          (median_scores.tmscore >= 0.2) & \
          (median_scores.fraction_binder >= 0.2) & \
          (median_scores.rmsd <= 15))
median_scores_filtered = median_scores[select]
```

In [ ]:
```
median_scores_filtered.shape
```

Out[ ]: (83, 7)

See how many clusters are left after filtering:

In [ ]:
```
median_scores_filtered.cluster.unique().shape
```

Out[ ]: (3,)

Function to format tables:

In [ ]:
```
import seaborn as sns
cm_r = sns.color_palette("mako_r", as_cmap=True)
cm = sns.color_palette("mako", as_cmap=True)
```

In [ ]:
```
def make_pretty(styler):
    styler.format(precision=2)
    styler.background_gradient(axis=0, cmap=cm_r, subset=pd.IndexSlice[:,"cl
    styler.background_gradient(axis=0, cmap=cm_r, subset=pd.IndexSlice[:,"tm
    styler.background_gradient(axis=0, cmap=cm, subset=pd.IndexSlice[:,"rmsd
    styler.background_gradient(axis=0, cmap=cm_r, subset=pd.IndexSlice[:,"fr
    return styler
```

### RESULT 1: See clusters ranked by RMSD

Finally, we can rank the clusters and see which ones have a good combination of low RMSD and large cluster size. These ones are the most likely to contain the true binder.

In [ ]:
```
# Select the rows with the minimum RMSD for each cluster
select = median_scores_filtered.groupby('cluster').rmsd.idxmin()
columns = ['cluster', 'tmscore', 'rmsd', 'cluster_size', 'fraction_binder']
median_scores_filtered.loc[select,columns].sort_values(by='rmsd').style.pipe
```

Out[ ]:

| | cluster | tmscore | rmsd | cluster_size | fraction_binder |
|---|---|---|---|---|---|
| **59** | Q7XL74.pdb_B | 0.90 | 1.49 | 40.00 | 0.94 |
| **22** | Q6Z1A9.pdb_B | 0.55 | 10.65 | 49.00 | 0.84 |
| **4** | P42211.pdb_B | 0.66 | 10.75 | 10.00 | 0.90 |

Here we can see that the cluster `Q7XL74.pdb_B` has the lowest median RMSD and the highest median TM-score. It also has a very large size with 40 members.

### RESULT 2: See clusters ranked by size

In [ ]:
```
# Select the rows with the minimum RMSD for each cluster
select = median_scores_filtered.groupby('cluster').rmsd.idxmin()
columns = ['cluster', 'tmscore', 'rmsd', 'cluster_size', 'fraction_binder']
median_scores_filtered.loc[select, columns].sort_values(by='cluster_size', a
```

Out[ ]:

| | cluster | tmscore | rmsd | cluster_size | fraction_binder |
|---|---|---|---|---|---|
| 22 | Q6Z1A9.pdb_B | 0.55 | 10.65 | 49.00 | 0.84 |
| 59 | Q7XL74.pdb_B | 0.90 | 1.49 | 40.00 | 0.94 |
| 4 | P42211.pdb_B | 0.66 | 10.75 | 10.00 | 0.90 |

Here we can see that the cluster `Q6Z1A9.pdb_B` is the largest cluster with 49 members, but it has a much worse RMSD than `Q7XL74.pdb_B`. It would be interesting to look at both and see how they compare. So we managed to reduce more than 43,000 starting models to only two promising clusters!

For this example we know that in our list of candidate binders there are 14 homologues of the true binder protein. We can find out which clusters contain these homologues:

In [ ]:

```
homologues = ['Q6YY33', 'Q6YY34', 'A0A0P0VKX7', 'Q0JCK8', 'Q6YY31', 'Q7XJV3'
         'A0A0N7KFK3', 'Q7XJV0', 'A0A0P0WB87', 'Q6EPT4', 'Q0J314', 'Q6EPT2',
         'Q8S5W0', 'Q2QSQ7']
```

In [ ]:

```
clusters[clusters.member.isin(homologues)][['complex','merged_rep','iptm','i
```

Out[ ]:

| | complex | merged_rep | iptm | iptm+ptm |
|---|---|---|---|---|
| 6 | 6Q76-1_Q6EPT2-1 | Q7XL74.pdb_B | 0.913843 | 0.847186 |
| 14 | 6Q76-1_Q6EPT4-1 | Q7XL74.pdb_B | 0.903790 | 0.819935 |
| 26 | 6Q76-1_Q2QSQ7-1 | Q7XL74.pdb_B | 0.892084 | 0.806683 |
| 30 | 6Q76-1_Q0J314-1 | Q7XL74.pdb_B | 0.886320 | 0.820720 |
| 38 | 6Q76-1_Q8S5W0-1 | Q7XL74.pdb_B | 0.875703 | 0.805274 |
| 43 | 6Q76-1_Q6YY34-1 | Q7XL74.pdb_B | 0.867315 | 0.839418 |
| 47 | 6Q76-1_Q7XJV3-1 | Q7XL74.pdb_B | 0.862152 | 0.831752 |
| 48 | 6Q76-1_Q6YY33-1 | Q7XL74.pdb_B | 0.860912 | 0.830552 |
| 55 | 6Q76-1_A0A0N7KFK3-1 | Q7XL74.pdb_B | 0.857097 | 0.819887 |
| 57 | 6Q76-1_A0A0P0VKX7-1 | Q7XL74.pdb_B | 0.854087 | 0.827703 |
| 63 | 6Q76-1_A0A0P0WB87-1 | Q7XL74.pdb_B | 0.843057 | 0.768634 |
| 71 | 6Q76-1_Q6YY31-1 | Q7XL74.pdb_B | 0.825141 | 0.805893 |
| 76 | 6Q76-1_Q0JCK8-1 | Q7XL74.pdb_B | 0.818324 | 0.793344 |
| 84 | 6Q76-1_Q7XJV0-1 | Q7XL74.pdb_B | 0.806434 | 0.778532 |

They are all in the top cluster by RMSD!

### 2. Make pymol sessions for the top clusters with `make_pymol_sessions`

Run the following command to select the top clusters that we saw above, make pymol sessions of the top clusters, and optionally do structural clustering on each cluster to find subclusters:

```
alphacrv-rank \
  --clusters_dir ./examples/AVRPia/AVRPia_vs_rice_clusters \
  --min_members 5 \
  --min_tmscore 0.2 \
  --max_rmsd 15 \
  --cluster_clusters
```

The program will show you the top clusters that will be used to make the pymol sessions. You can press `Enter` to continue, or exit the program with `Ctrl+C` to change the filtering parameters.

```
(env) AlphaCRV$ alphacrv-rank \
--clusters_dir ./examples/AVRPia/AVRPia_vs_rice_clusters \
--min_members 5 \
--min_tmscore 0.2 \
--max_rmsd 15 \
--cluster_clusters
INFO:root:Identified 3 top clusters.
INFO:root:Top clusters:

        cluster  tmscore    rmsd  cluster_size
fraction_binder
0  Q6Z1A9.pdb_B  0.54957  10.655          49.0
0.839820
1  Q7XL74.pdb_B  0.89660   1.490          40.0
0.935897
2  P42211.pdb_B  0.66389  10.750          10.0
0.902703

Press Enter to continue, or Ctrl+C to exit and select
different filtering parameters:
INFO:root:Copying pdbs from the top clusters...
INFO:root:Making Pymol sessions...
PyMOL>select chain B AND model P42211_repB
 Selector: selection "sele" defined with 6294 atoms.
PyMOL>bg white
PyMOL>set ray_shadow, 0
 Setting: ray_shadow set to off.
PyMOL>color grey80
 Executive: Colored 72547 atoms.
PyMOL>select chain A
 Selector: selection "sele" defined with 10530 atoms.
PyMOL>color slate, sele
 Executive: Colored 10530 atoms.
PyMOL>delete all
PyMOL>select chain B AND model Q6Z1A9_repB
 Selector: selection "sele" defined with 3855 atoms.
PyMOL>bg white
PyMOL>set ray_shadow, 0
 Setting: ray_shadow set to off.
PyMOL>color grey80
 Executive: Colored 368033 atoms.
PyMOL>select chain A
 Selector: selection "sele" defined with 51597 atoms.
PyMOL>color slate, sele
 Executive: Colored 51597 atoms.
PyMOL>delete all
PyMOL>select chain B AND model Q7XL74_repB
 Selector: selection "sele" defined with 1365 atoms.
PyMOL>bg white
PyMOL>set ray_shadow, 0
 Setting: ray_shadow set to off.
```

```
PyMOL>color grey80
 Executive: Colored 92930 atoms.
PyMOL>select chain A
 Selector: selection "sele" defined with 42120 atoms.
PyMOL>color slate, sele
 Executive: Colored 42120 atoms.
PyMOL>delete all
INFO:root:Clustering clusters...
INFO:root:Clustering P42211.pdb_B
INFO:root:Running structural clustering...
INFO:root:Processing output...
INFO:root:Clustering Q6Z1A9.pdb_B
INFO:root:Running structural clustering...
INFO:root:Processing output...
INFO:root:Clustering Q7XL74.pdb_B
INFO:root:Running structural clustering...
INFO:root:Processing output...
INFO:root:Done!!
```

This command should create the following files in the `./examples/AVRPia/AVRPia_vs_rice_clusters/merged_clusters/` / `merged_clusters` directory:

- `clustered_clusters.csv` : Contains the subclusters for each of the top clusters.
- `cluster_<cluster_ID>/` : Contains the PDBs of each cluster, and a PyMol session with the cluster members.
- `cluster_<cluster_ID>_clusters/` : Contains the results of the `foldseek easy-cluster` run on the cluster members.

### Read clustered clusters

The following DataFrame contains the subclusters for each of the top clusters:

In [ ]:
```python
clustered_clusters = pd.read_csv(results_dir / 'merged_clusters/clustered_cl
```

In [ ]:
```python
clustered_clusters.head()
```

Out[ ]:

| | subcluster_rep | member | cluster |
|---|---|---|---|
| **0** | P42211.pdb_B | P42211 | P42211.pdb_B |
| **1** | P42211.pdb_B | Q6F4N5 | P42211.pdb_B |
| **2** | P42211.pdb_B | A0A0P0VAB8 | P42211.pdb_B |
| **3** | P42211.pdb_B | Q0D5V1 | P42211.pdb_B |
| **4** | P42211.pdb_B | Q8LNM1 | P42211.pdb_B |

Now we can look at the most interesting clusters and their subclusters in more detail:

### RESULT 1: Cluster Q7XL74.pdb_B (contains true binder homologs, top cluster by size)

In [ ]:
```
cluster = 'Q7XL74.pdb_B'
```

See subclusters:

In [ ]:
```
clustered_clusters[clustered_clusters.cluster==cluster]
```

Out[ ]:

| | subcluster_rep | member | cluster |
|---|---|---|---|
| 59 | A0A0P0UZA6.pdb_B | A0A0P0UZA6 | Q7XL74.pdb_B |
| 60 | Q7XL74.pdb_B | Q7XL74 | Q7XL74.pdb_B |
| 61 | Q7XL74.pdb_B | Q0IYI4 | Q7XL74.pdb_B |
| 62 | Q7XL74.pdb_B | Q75J02 | Q7XL74.pdb_B |
| 63 | Q7XL74.pdb_B | A0A0P0WEB4 | Q7XL74.pdb_B |
| 64 | Q7XL74.pdb_B | A0A0P0VRA1 | Q7XL74.pdb_B |
| 65 | Q7XL74.pdb_B | Q0JBG2 | Q7XL74.pdb_B |
| 66 | Q7XL74.pdb_B | A0A0P0XD00 | Q7XL74.pdb_B |
| 67 | Q7XL74.pdb_B | A0A0P0V126 | Q7XL74.pdb_B |
| 68 | Q7XL74.pdb_B | A3AYA2 | Q7XL74.pdb_B |
| 69 | Q7XL74.pdb_B | Q10KH9 | Q7XL74.pdb_B |
| 70 | Q7XL74.pdb_B | Q84TS2 | Q7XL74.pdb_B |
| 71 | Q7XL74.pdb_B | Q8LNN6 | Q7XL74.pdb_B |
| 72 | Q7XL74.pdb_B | Q8S181 | Q7XL74.pdb_B |
| 73 | Q7XL74.pdb_B | Q7G2B2 | Q7XL74.pdb_B |
| 74 | Q7XL74.pdb_B | A0A0P0XFR7 | Q7XL74.pdb_B |
| 75 | Q7XL74.pdb_B | A3ADD6 | Q7XL74.pdb_B |
| 76 | Q7XL74.pdb_B | Q6I5G4 | Q7XL74.pdb_B |
| 77 | Q7XL74.pdb_B | Q6YY31 | Q7XL74.pdb_B |
| 78 | Q7XL74.pdb_B | Q84TB9 | Q7XL74.pdb_B |
| 79 | Q7XL74.pdb_B | Q8LN41 | Q7XL74.pdb_B |
| 80 | Q7XL74.pdb_B | B7E663 | Q7XL74.pdb_B |
| 81 | Q7XL74.pdb_B | A0A0N7KFK3 | Q7XL74.pdb_B |
| 82 | Q7XL74.pdb_B | A0A0P0VKX7 | Q7XL74.pdb_B |
| 83 | Q7XL74.pdb_B | A0A0P0WB87 | Q7XL74.pdb_B |
| 84 | Q7XL74.pdb_B | Q0JCK8 | Q7XL74.pdb_B |
| 85 | Q7XL74.pdb_B | Q6EPT2 | Q7XL74.pdb_B |
| 86 | Q7XL74.pdb_B | Q6YY33 | Q7XL74.pdb_B |
| 87 | Q7XL74.pdb_B | Q6YY34 | Q7XL74.pdb_B |
| 88 | Q7XL74.pdb_B | Q7XJV0 | Q7XL74.pdb_B |
| 89 | Q7XL74.pdb_B | Q7XJV3 | Q7XL74.pdb_B |
| 90 | Q7XL74.pdb_B | Q2QSQ7 | Q7XL74.pdb_B |
| 91 | Q7XL74.pdb_B | A0A0P0V0T5 | Q7XL74.pdb_B |
| 92 | Q7XL74.pdb_B | A2ZV04 | Q7XL74.pdb_B |

| | subcluster_rep | member | cluster |
|---|---|---|---|
| **93** | Q7XL74.pdb_B | Q5JL91 | Q7XL74.pdb_B |
| **94** | Q7XL74.pdb_B | Q8S5W0 | Q7XL74.pdb_B |
| **95** | Q7XL74.pdb_B | A0A0P0Y5Q0 | Q7XL74.pdb_B |
| **96** | Q7XL74.pdb_B | Q0J314 | Q7XL74.pdb_B |
| **97** | Q7XL74.pdb_B | Q6EPT4 | Q7XL74.pdb_B |
| **98** | Q7XL74.pdb_B | Q8LMS1 | Q7XL74.pdb_B |

Here we have two subclusters ( `subcluster_rep` column). However, one of them ( `A0A0P0UZA6.pdb_B` ) has only one member, and if we look at it in the PyMOL session we can see that it is very similar to the other complexes. The only difference is the HMA domain being shorter. So for our purposes it would be ok to take them all together.

We can see in the `median_scores` DataFrame the model that has the best alignments to all other structures (best representative of the cluster).

In [ ]:
```
median_scores[median_scores.cluster==cluster].sort_values(by='rmsd').head(10
```

Out[ ]:

| | cluster | member | tmscore | rmsd | aligned_length | cluster_size | fraction_binde |
|---|---|---|---|---|---|---|---|
| **59** | Q7XL74.pdb_B | A0A0N7KFK3 | 0.89660 | 1.49 | 141.0 | 40.0 | 0.93589 |
| **61** | Q7XL74.pdb_B | A0A0P0V0T5 | 0.89782 | 1.50 | 140.0 | 40.0 | 0.96000 |
| **93** | Q7XL74.pdb_B | Q84TS2 | 0.90749 | 1.52 | 140.0 | 40.0 | 0.97297 |
| **62** | Q7XL74.pdb_B | A0A0P0V126 | 0.90529 | 1.58 | 140.0 | 40.0 | 0.97297 |
| **90** | Q7XL74.pdb_B | Q7XJV3 | 0.90317 | 1.62 | 141.0 | 40.0 | 0.94805 |
| **84** | Q7XL74.pdb_B | Q6YY31 | 0.90490 | 1.62 | 141.0 | 40.0 | 0.94805 |
| **63** | Q7XL74.pdb_B | A0A0P0VKX7 | 0.87889 | 1.74 | 142.0 | 40.0 | 0.93670 |
| **85** | Q7XL74.pdb_B | Q6YY33 | 0.88542 | 1.74 | 142.0 | 40.0 | 0.94871 |
| **70** | Q7XL74.pdb_B | A2ZV04 | 0.89069 | 1.74 | 138.0 | 40.0 | 0.97222 |
| **86** | Q7XL74.pdb_B | Q6YY34 | 0.88160 | 1.82 | 142.0 | 40.0 | 0.93670 |

We can also select the IDs of the binder proteins in this cluster, and run them through a tool such as DAVID to perform enrichment analysis.

In [ ]:
```
members = clusters[clusters.merged_rep==cluster].member
```

In [ ]:
```
for m in members:
    print(m)
```

```
A0A0P0UZA6
Q6EPT2
Q8LN41
Q6EPT4
Q6I5G4
Q7XL74
Q84TB9
Q2QSQ7
A0A0P0WEB4
Q0J314
A2ZV04
Q5JL91
Q8S5W0
Q8LNN6
Q75J02
Q6YY34
B7E663
Q7XJV3
Q6YY33
Q8LMS1
A0A0P0XD00
Q7G2B2
A0A0N7KFK3
A0A0P0V0T5
A0A0P0VKX7
A0A0P0XFR7
A0A0P0Y5Q0
A0A0P0WB87
Q10KH9
A0A0P0V126
A3ADD6
Q6YY31
A3AYA2
Q0JBG2
Q0JCK8
Q8S181
A0A0P0VRA1
Q7XJV0
Q0IYI4
Q84TS2
```

Looking at the `examples/AVRPia/AVRPia_vs_rice_clusters/merged_clusters/cluster_Q7XL74.pdb_B/session.pse` file in PyMol, we can visualize the cluster and produce a figure like this:

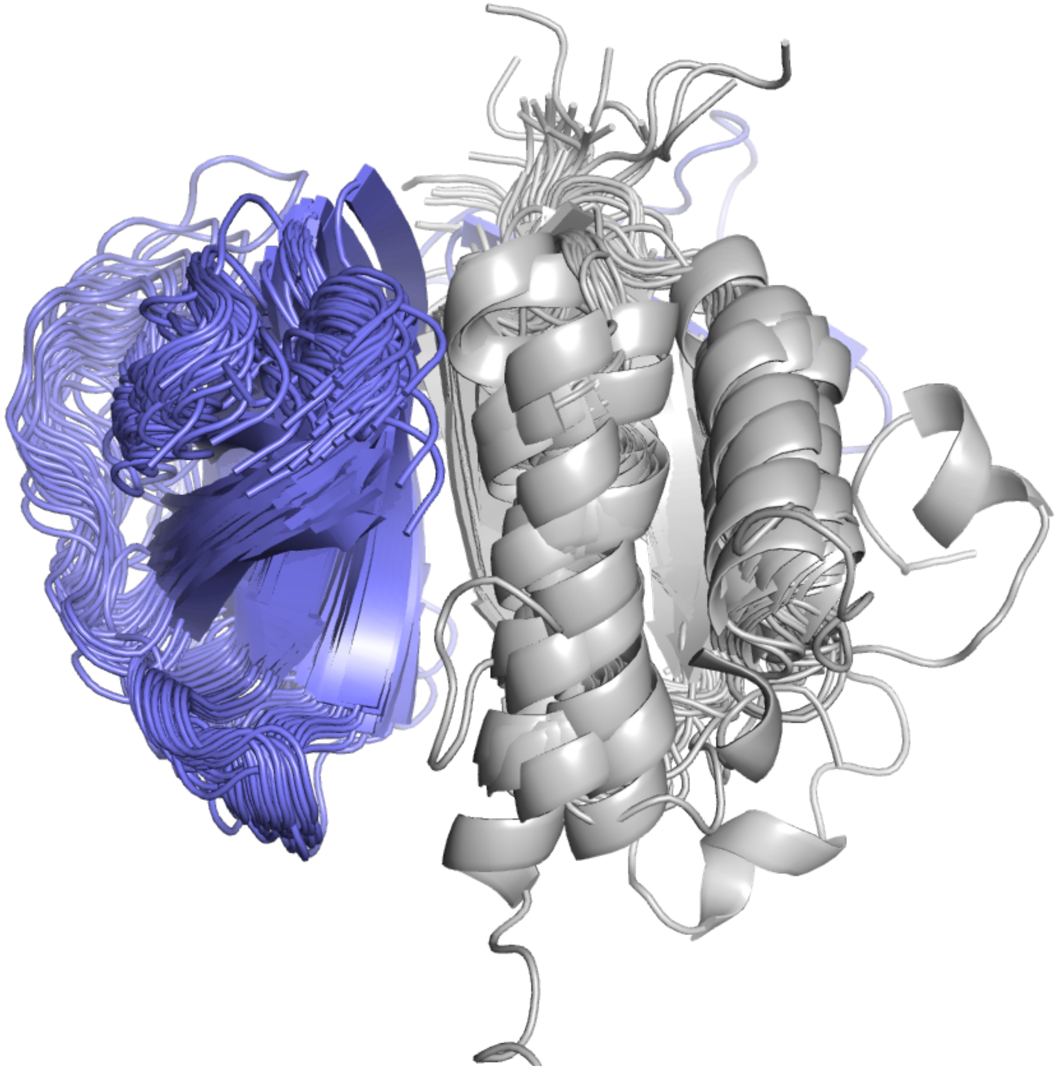

When aligning the HMA domains of the binder structures (grey), the binding of the AVR-Pia protein seems pretty consistent across most of the models in the cluster.

Tip: In PyMol, you can remove the residues with low pLDDT score to make the figure cleaner, with the command:

```
select b < 50; remove sele
```

### Clustering AVR-Pik

This notebook will guide you through the process of clustering 65 complexes of the AVR-Pik protein against a sample of rice proteins. All of the models are dimeric. The 65 models originate from a proteome-wide screen of AVR-Pik against the rice proteome (O. sativa subsp. japonica, 43,000+ initial models). To select this sample we first ran AlphaCRV on all the models, and the 65 structures are part of the best clusters that were identified. Running the pipeline on these models should be enough to reproduce the results.

### Prerequisites

- Install AlphaCRV on a conda environment and activate it
- Download the sequences, models and results for this example at Zenodo

### 1. Cluster the models with the `clustering` command

Run the following command to cluster the models. Make sure to change the paths according to your system:

```
alphacrv-cluster \
  --bait ./examples/AVRPik/AVRPik.fasta \
  --binders ./examples/AVRPik/AVRPik_binders.fasta \
  --models_dir ./examples/AVRPik/AVRPik_vs_rice_models \
  --destination ./examples/AVRPik/AVRPik_vs_rice_clusters \
  --cpus 8
```

After collecting the quality scores from the models in `--models_dir`, it will count how many models are there with an ipTM scores higher than the threshold (0.75 by default). It will prompt you to confirm or to modify the threshold. After that, it will proceed with the clustering.

The full output is as follows:

```
(env) AlphaCRV$ alphacrv-cluster \
--bait ./examples/AVRPik/AVRPik.fasta \
--binders ./examples/AVRPik/AVRPik_binders.fasta \
--models_dir ./examples/AVRPik/AVRPik_vs_rice_models \
--destination ./examples/AVRPik/AVRPik_vs_rice_clusters \
--cpus 8
INFO:root:Getting quality scores for models in
examples/AVRPik/AVRPik_vs_rice_models...
INFO:root:Found 65 model directories with quality scores.
Will select 65 models with iptm >= 0.75. Press enter to
continue, or enter a new threshold:
INFO:root:Trimming binder molecules to keep only regions with
an average PAE against the bait of up to 10.0...
INFO:root:Processed 65 complexes.
INFO:root:Writing trimmed sequences to fasta file.
INFO:root:Running sequence clustering...
INFO:root:Processing output...
INFO:root:Running structural clustering...
INFO:root:Processing output...
INFO:root:Aligning all vs all members of each cluster...
INFO:root:Aligning cluster 1 of 4...
INFO:root:11 members.
INFO:root:Aligning cluster 2 of 4...
INFO:root:21 members.
INFO:root:Aligning cluster 3 of 4...
INFO:root:13 members.
INFO:root:Aligning cluster 4 of 4...
INFO:root:20 members.
INFO:root:Calculating median alignment scores...
INFO:root:Done!!
```

After this step you will have a directory with the following structure:

```
(env) AlphaCRV$ ll examples/AVRPik/AVRPik_vs_rice_clusters/
total 24K
-rw-r--r-- 1 example g-example 1.3K Jan  3 16:30
binders_regions.csv
drwxr-xr-x 1 example g-example    0 Jan  3 16:31
merged_clusters/
drwxr-xr-x 1 example g-example    0 Jan  3 16:30
pdbs_trimmed/
drwxr-xr-x 1 example g-example    0 Jan  3 16:30 seqclusters/
drwxr-xr-x 1 example g-example    0 Jan  3 16:30 strclusters/
-rw-r--r-- 1 example g-example  19K Jan  3 16:30
trimmed_binders.fasta
```

Now let's look at some of the important files:

In [ ]:
```
from pathlib import Path
import pandas as pd
```

In [ ]:
```
results_dir = Path('./AVRPik/AVRPik_vs_rice_clusters/')
```

### See clusters

The `merged_clusters.csv` file contains the list of models with their corresponding sequence, structure and merged clusters. It also has the quality scores provided by AlphaFold.

In [ ]:
```
clusters = pd.read_csv(results_dir / 'merged_clusters/merged_clusters.csv')
```

In [ ]:
```
clusters
```

Out[ ]:

| | complex | str_rep | seq_rep | merged_rep | meml |
|---|---|---|---|---|---|
| **0** | 6R8K-1_A0A0P0Y5A4-1 | Q67VV7.pdb_B | Q6ZEZ7 | Q0DEU2.pdb_B | A0A0P0Y5 |
| **1** | 6R8K-1_A0A0P0YB11-1 | Q67VV7.pdb_B | Q6ZEZ7 | Q0DEU2.pdb_B | A0A0P0YI |
| **2** | 6R8K-1_A0A0P0VEM6-1 | Q67VV7.pdb_B | Q6ZEZ7 | Q0DEU2.pdb_B | A0A0P0VE |
| **3** | 6R8K-1_Q8L3T8-1 | Q67VV7.pdb_B | Q6ZEZ7 | Q0DEU2.pdb_B | Q8L3 |
| **4** | 6R8K-1_Q2RAL3-1 | Q6K9R5.pdb_B | Q6ZBC3 | Q6K9R5.pdb_B | Q2RA |
| **...** | ... | ... | ... | ... | |
| **60** | 6R8K-1_Q0DBF4-1 | A3BDZ2.pdb_B | Q0DBF4 | A3BDZ2.pdb_B | Q0DE |
| **61** | 6R8K-1_A3ADD6-1 | Q6K9R5.pdb_B | Q7G2B2 | Q6K9R5.pdb_B | A3AD |
| **62** | 6R8K-1_Q0JCK8-1 | Q6K9R5.pdb_B | A0A0N7KFK3 | Q6K9R5.pdb_B | Q0JC |
| **63** | 6R8K-1_Q0JKW0-1 | A0A0P0WKJ4.pdb_B | A0A0P0Y219 | A0A0P0WKJ4.pdb_B | Q0JK |
| **64** | 6R8K-1_A3BDZ2-1 | A3BDZ2.pdb_B | A3BDZ2 | A3BDZ2.pdb_B | A3BD |

65 rows × 7 columns

The columns of the `clusters` DataFrame are:

- `complex` : The name of the complex. This is the same name as the directory where the model is stored.
- `str_rep` : Name of the structure cluster representative
- `seq_rep` : Name of the sequence cluster representative
- `merged_rep` : Name of the merged cluster representative (sequence + structure)
- `member` : The ID of the binder protein
- `iptm` : The ipTM score of the model
- `iptm+ptm` : The ipTM+PTM score of the model (it is calculated by AlphaFold as $0.8 ipTM + 0.2$pTM)

See the number of different merged clusters:

In [ ]:
```
clusters.merged_rep.unique().shape
```

Out[ ]: (4,)

For this example, the models and sequences from the 65 binder proteins were summarized in 4 clusters. Much fewer structures to sort through!

### See alignment scores

Alignment scores are calcualted for each cluster by aligning all vs all members of the cluster.

In [ ]:
```
alignment_scores = pd.read_csv(results_dir / 'merged_clusters/alignment_scor
```

In [ ]:
```
alignment_scores.head()
```

Out[ ]:

| | cluster | ref | member | tmscore_ref | tmscore_m | aligned_length | rms |
|---|---|---|---|---|---|---|---|
| **0** | Q0DEU2.pdb_B | A0A0P0Y5A4 | A0A0P0YB11 | 0.88555 | 0.97405 | 432 | 1.4 |
| **1** | Q0DEU2.pdb_B | A0A0P0Y5A4 | A0A0P0VEM6 | 0.86424 | 0.91255 | 440 | 3.0 |
| **2** | Q0DEU2.pdb_B | A0A0P0Y5A4 | Q8L3T8 | 0.85603 | 0.93629 | 430 | 2.5 |
| **3** | Q0DEU2.pdb_B | A0A0P0Y5A4 | B9EWP2 | 0.86538 | 0.95866 | 425 | 2.1 |
| **4** | Q0DEU2.pdb_B | A0A0P0Y5A4 | Q7XV05 | 0.37127 | 0.61033 | 271 | 13.[illegible] |

The columns of the `alignment_scores` DataFrame are:

- `cluster` : The name of the cluster
- `ref` : Binder ID of the reference structure in the alignment
- `member` : Binder ID of the second structure in the alignment
- `tmscore_ref` : TM-score based on the reference structure
- `tmscore_m` : TM-score based on the second structure
- `aligned_length` : Length of the alignment
- `rmsd` : RMSD of the alignment

Based on these scores, the median scores are calculated for each cluster member to find the best representative of the cluster (the one with lowest RMSD score to the other members).

### Read median scores and find top clusters

Now we can rank the clusters based on the median alignment scores of the cluster representatives:

In [ ]:
```
median_scores = pd.read_csv(results_dir / 'merged_clusters/median_scores.csv
```

In [ ]:
```
median_scores.shape
```

Out[ ]:
```
(65, 7)
```

The `median_scores` DataFrame contains the median alignment scores of each cluster member when aligned to all other members of the same cluster.

In [ ]:
```
median_scores.head()
```

Out[ ]:

| | cluster | member | tmscore | rmsd | aligned_length | cluster_size | fraction_[illegible] |
|---|---|---|---|---|---|---|---|
| 0 | A0A0P0WKJ4.pdb_B | A0A0P0W913 | 0.43642 | 15.29 | 250.0 | 20.0 | 0.8[illegible] |
| 1 | A0A0P0WKJ4.pdb_B | A0A0P0WKJ4 | 0.31741 | 13.10 | 219.0 | 20.0 | 0.5[illegible] |
| 2 | A0A0P0WKJ4.pdb_B | A0A0P0WQF2 | 0.40264 | 12.55 | 226.0 | 20.0 | 0.7[illegible] |
| 3 | A0A0P0WKJ4.pdb_B | A0A0P0XBZ1 | 0.26659 | 17.31 | 266.0 | 20.0 | 0.4[illegible] |
| 4 | A0A0P0WKJ4.pdb_B | A0A0P0XUC4 | 0.31355 | 16.23 | 268.0 | 20.0 | 0.[illegible] |

The columns of the `median_scores` DataFrame are:

- `cluster` : The name of the cluster
- `member` : ID of the cluster member (binder molecule)
- `tmscore` : Median TM-score of the complex against all other complexes in this cluster
- `rmsd` : Median RMSD of the complex against all other complexes in this cluster
- `aligned_length` : Median length of the alignment
- `cluster_size`
- `fraction_binder` : In average, how much of the binder molecule is included in the alignments of this complex against all other complexes (calculated as `(aligned_length - bait_length)/binder_length` ). This is just meant to be an approximation of how complete the aligmnents are for this cluster member.

The next step is to select the cluster representatives. For this, we first need to filter out the cluster members with poor quality alignments, according to the following criteria:

- Small size
- Low median TM-score
- High median RMSD
- Low fraction of the binder aligned in the cluster representative

In [ ]:
```
# Select the clusters with the following criteria:
select = ((median_scores.cluster_size >= 5) & \
            (median_scores.tmscore >= 0.2) & \
            (median_scores.fraction_binder >= 0.2) & \
            (median_scores.rmsd <= 15))
median_scores_filtered = median_scores[select]
```

In [ ]:
```
median_scores_filtered.shape
```

Out[ ]:
```
(51, 7)
```

See how many clusters are left after filtering:

In [ ]:
```
median_scores_filtered.cluster.unique().shape
```

Out[ ]:
```
(4,)
```

Function to format tables:

In [ ]:
```
import seaborn as sns
cm_r = sns.color_palette("mako_r", as_cmap=True)
cm = sns.color_palette("mako", as_cmap=True)
```

In [ ]:
```
def make_pretty(styler):
    styler.format(precision=2)
    styler.background_gradient(axis=0, cmap=cm_r, subset=pd.IndexSlice[:,"cl
    styler.background_gradient(axis=0, cmap=cm_r, subset=pd.IndexSlice[:,"tm
    styler.background_gradient(axis=0, cmap=cm, subset=pd.IndexSlice[:,"rmsd
    styler.background_gradient(axis=0, cmap=cm_r, subset=pd.IndexSlice[:,"fr
    return styler
```

### RESULT 1: See clusters ranked by RMSD

Finally, we can rank the clusters and see which ones have a good combination of low RMSD and large cluster size. These ones are the most likely to contain the true binder.

In [ ]:
```
# Select the rows with the minimum RMSD for each cluster
select = median_scores_filtered.groupby('cluster').rmsd.idxmin()
columns = ['cluster', 'tmscore', 'rmsd', 'cluster_size', 'fraction_binder']
median_scores_filtered.loc[select,columns].sort_values(by='rmsd').style.pipe
```

Out[ ]:

| | cluster | tmscore | rmsd | cluster_size | fraction_binder |
|---|---|---|---|---|---|
| 41 | Q0DEU2.pdb_B | 0.94 | 2.17 | 11.00 | 0.99 |
| 28 | A3BDZ2.pdb_B | 0.82 | 3.38 | 13.00 | 0.94 |
| 54 | Q6K9R5.pdb_B | 0.31 | 4.99 | 21.00 | 0.31 |
| 9 | A0A0P0WKJ4.pdb_B | 0.42 | 10.93 | 20.00 | 0.86 |

Here we can see that the cluster `Q0DEU2.pdb_B` has the lowest median RMSD and the highest median TM-score. However, it has a relatively small size with only 11 members.

### RESULT 2: See clusters ranked by size

In [ ]:
```
# Select the rows with the minimum RMSD for each cluster
select = median_scores_filtered.groupby('cluster').rmsd.idxmin()
columns = ['cluster', 'tmscore', 'rmsd', 'cluster_size', 'fraction_binder']
median_scores_filtered.loc[select, columns].sort_values(by='cluster_size', a
```

Out[ ]:

| | cluster | tmscore | rmsd | cluster_size | fraction_binder |
|---|---|---|---|---|---|
| **54** | Q6K9R5.pdb_B | 0.31 | 4.99 | 21.00 | 0.31 |
| **9** | A0A0P0WKJ4.pdb_B | 0.42 | 10.93 | 20.00 | 0.86 |
| **28** | A3BDZ2.pdb_B | 0.82 | 3.38 | 13.00 | 0.94 |
| **41** | Q0DEU2.pdb_B | 0.94 | 2.17 | 11.00 | 0.99 |

Here we can see that the cluster `Q6K9R5.pdb_B` is the largest cluster with 21 members. It also has a low RMSD of 4.99, although the median TM-score is not the highest. This cluster, along with `A3BDZ2.pdb_B` and `A3BDZ2.pdb_B` would be good candidates for further analysis. Only three clusters with a total of 45 structures, from more than 43,000 starting models!

For this example we know that in our list of candidate binders there are 6 homologues of the true binder protein. We can find out which clusters contain these homologues:

In [ ]:
```
homologues = ['Q7XJV3', 'Q6EPT2','A0A0N7KFK3', 'Q6YY31', 'Q6EPT4', 'Q0JCK8']
```

In [ ]:
```
clusters[clusters.member.isin(homologues)][['complex','merged_rep','iptm','i
```

Out[ ]:

| | complex | merged_rep | iptm | iptm+ptm |
|---|---|---|---|---|
| **14** | 6R8K-1_Q7XJV3-1 | Q6K9R5.pdb_B | 0.828985 | 0.807492 |
| **33** | 6R8K-1_Q6EPT2-1 | Q6K9R5.pdb_B | 0.806888 | 0.763640 |
| **36** | 6R8K-1_A0A0N7KFK3-1 | Q6K9R5.pdb_B | 0.800443 | 0.782608 |
| **45** | 6R8K-1_Q6YY31-1 | Q6K9R5.pdb_B | 0.784981 | 0.775073 |
| **56** | 6R8K-1_Q6EPT4-1 | Q6K9R5.pdb_B | 0.766761 | 0.710590 |
| **62** | 6R8K-1_Q0JCK8-1 | Q6K9R5.pdb_B | 0.756162 | 0.744660 |

They are all in the largest cluster!

### 2. Make pymol sessions for the top clusters with `make_pymol_sessions`

Run the following command to select the top clusters that we saw above, make pymol sessions of the top clusters, and optionally do structural clustering on each cluster to find subclusters:

```
alphacrv-rank \
  --clusters_dir /path/to/destination/6R8K_clusters \
  --min_members 5 \
  --min_tmscore 0.2 \
  --max_rmsd 15 \
  --cluster_clusters
```

The program will show you the top clusters that will be used to make the pymol sessions. You can press `Enter` to continue, or exit the program with `Ctrl+C` to change the filtering parameters.

```
(env) AlphaCRV$ alphacrv-rank \
--clusters_dir ./examples/AVRPik/AVRPik_vs_rice_clusters \
--min_members 5 \
--min_tmscore 0.2 \
--max_rmsd 15 \
--cluster_clusters
INFO:root:Identified 4 top clusters.
INFO:root:Top clusters:

            cluster   tmscore    rmsd  cluster_size
fraction_binder
0      Q6K9R5.pdb_B  0.313905   4.985          21.0
0.305328
1  A0A0P0WKJ4.pdb_B  0.421400  10.930          20.0
0.856436
2      A3BDZ2.pdb_B  0.824705   3.385          13.0
0.944444
3      Q0DEU2.pdb_B  0.940945   2.165          11.0
0.987633

Press Enter to continue, or Ctrl+C to exit and select
different filtering parameters:
INFO:root:Copying pdbs from the top clusters...
INFO:root:Making Pymol sessions...
PyMOL>select chain B AND model A0A0P0WKJ4_repB
 Selector: selection "sele" defined with 3937 atoms.
PyMOL>bg white

...
PyMOL>select chain B AND model Q6K9R5_repB
 Selector: selection "sele" defined with 1397 atoms.
PyMOL>bg white
PyMOL>set ray_shadow, 0
 Setting: ray_shadow set to off.
PyMOL>color grey80
 Executive: Colored 61825 atoms.
PyMOL>select chain A
 Selector: selection "sele" defined with 31164 atoms.
PyMOL>color slate, sele
 Executive: Colored 31164 atoms.
PyMOL>delete all
INFO:root:Clustering clusters...
INFO:root:Clustering A0A0P0WKJ4.pdb_B
INFO:root:Running structural clustering...
INFO:root:Processing output...
INFO:root:Clustering A3BDZ2.pdb_B
INFO:root:Running structural clustering...
INFO:root:Processing output...
INFO:root:Clustering Q0DEU2.pdb_B
INFO:root:Running structural clustering...
INFO:root:Processing output...
```

```
INFO:root:Clustering Q6K9R5.pdb_B
INFO:root:Running structural clustering...
INFO:root:Processing output...
INFO:root:Done!!
```

This command should create the following files in the `--clusters_dir` / `merged_clusters` directory:

- `clustered_clusters.csv` : Contains the subclusters for each of the top clusters.
- `cluster_<cluster_ID>/` : Contains the PDBs of each cluster, and a PyMol session with the cluster members.
- `cluster_<cluster_ID>_clusters/` : Contains the results of the `foldseek easy-cluster` run on the cluster members.

#### Read clustered clusters

The following DataFrame contains the subclusters for each of the top clusters:

In [ ]:
```
clustered_clusters = pd.read_csv(results_dir / 'merged_clusters/clustered_cl
```

In [ ]:
```
clustered_clusters.head()
```

Out[ ]:

| | subcluster_rep | member | cluster |
|---|---|---|---|
| **0** | A0A0P0W913.pdb_B | A0A0P0W913 | A0A0P0WKJ4.pdb_B |
| **1** | A0A0P0W913.pdb_B | A0A0P0WQF2 | A0A0P0WKJ4.pdb_B |
| **2** | A0A0P0WKJ4.pdb_B | A0A0P0WKJ4 | A0A0P0WKJ4.pdb_B |
| **3** | A0A0P0WKJ4.pdb_B | Q0JKW0 | A0A0P0WKJ4.pdb_B |
| **4** | A0A0P0XBZ1.pdb_B | A0A0P0XBZ1 | A0A0P0WKJ4.pdb_B |

Now we can look at the most interesting clusters and their subclusters in more detail:

#### RESULT 1: Cluster Q6K9R5.pdb_B (contains true binder homologs, top cluster by size)

In [ ]:
```
cluster = 'Q6K9R5.pdb_B'
```

See subclusters:

In [ ]:
```
clustered_clusters[clustered_clusters.cluster==cluster]
```

Out[ ]:

| | subcluster_rep | member | cluster |
|---|---|---|---|
| **44** | Q6K9R5.pdb_B | Q6K9R5 | Q6K9R5.pdb_B |
| **45** | Q6K9R5.pdb_B | A0A0N7KFK3 | Q6K9R5.pdb_B |
| **46** | Q6K9R5.pdb_B | B7E663 | Q6K9R5.pdb_B |
| **47** | Q6K9R5.pdb_B | Q6EPT2 | Q6K9R5.pdb_B |
| **48** | Q6K9R5.pdb_B | Q7XJV3 | Q6K9R5.pdb_B |
| **49** | Q6K9R5.pdb_B | Q6YY31 | Q6K9R5.pdb_B |
| **50** | Q6K9R5.pdb_B | Q7G2B2 | Q6K9R5.pdb_B |
| **51** | Q6K9R5.pdb_B | A3ADD6 | Q6K9R5.pdb_B |
| **52** | Q6K9R5.pdb_B | Q94CS5 | Q6K9R5.pdb_B |
| **53** | Q6K9R5.pdb_B | Q0JCK8 | Q6K9R5.pdb_B |
| **54** | Q6K9R5.pdb_B | Q6ZBC3 | Q6K9R5.pdb_B |
| **55** | Q6K9R5.pdb_B | Q2RAL3 | Q6K9R5.pdb_B |
| **56** | Q6K9R5.pdb_B | Q5JL91 | Q6K9R5.pdb_B |
| **57** | Q6K9R5.pdb_B | Q84TB9 | Q6K9R5.pdb_B |
| **58** | Q6K9R5.pdb_B | A0A0P0XYR1 | Q6K9R5.pdb_B |
| **59** | Q6K9R5.pdb_B | Q10N90 | Q6K9R5.pdb_B |
| **60** | Q6K9R5.pdb_B | Q2QZ01 | Q6K9R5.pdb_B |
| **61** | Q6K9R5.pdb_B | Q7EY69 | Q6K9R5.pdb_B |
| **62** | Q6K9R5.pdb_B | Q6EPT4 | Q6K9R5.pdb_B |
| **63** | Q6K9R5.pdb_B | Q8L4I2 | Q6K9R5.pdb_B |
| **64** | Q6K9R5.pdb_B | Q8LJL3 | Q6K9R5.pdb_B |

Only one subcluster ( `subcluster_rep` column).

We can see in the `median_scores` DataFrame the model that has the best alignments to all other structures (best representative of the cluster).

In [ ]:

```
median_scores[median_scores.cluster==cluster].sort_values(by='rmsd').head(10
```

Out[ ]:

| | cluster | member | tmscore | rmsd | aligned_length | cluster_size | fraction_binc |
|---|---|---|---|---|---|---|---|
| **54** | Q6K9R5.pdb_B | Q6EPT4 | 0.313905 | 4.985 | 167.5 | 21.0 | 0.3053 |
| **51** | Q6K9R5.pdb_B | Q2RAL3 | 0.772890 | 5.370 | 163.0 | 21.0 | 0.9722 |
| **57** | Q6K9R5.pdb_B | Q6ZBC3 | 0.591925 | 5.620 | 161.0 | 21.0 | 0.9315 |
| **61** | Q6K9R5.pdb_B | Q84TB9 | 0.608545 | 6.680 | 162.0 | 21.0 | 0.9718 |
| **45** | Q6K9R5.pdb_B | A0A0P0XYR1 | 0.548765 | 7.355 | 142.0 | 21.0 | 0.5903 |
| **50** | Q6K9R5.pdb_B | Q2QZ01 | 0.545625 | 7.420 | 142.0 | 21.0 | 0.5903 |
| **53** | Q6K9R5.pdb_B | Q6EPT2 | 0.575325 | 7.915 | 167.0 | 21.0 | 0.8915 |
| **58** | Q6K9R5.pdb_B | Q7EY69 | 0.544055 | 8.060 | 132.0 | 21.0 | 0.4698 |
| **49** | Q6K9R5.pdb_B | Q10N90 | 0.553010 | 8.125 | 139.0 | 21.0 | 0.5542 |
| **64** | Q6K9R5.pdb_B | Q94CS5 | 0.604640 | 8.180 | 163.0 | 21.0 | 0.9722 |

We can also select the IDs of the binder proteins in this cluster, and run them through a tool such as DAVID to perform enrichment analysis.

In [ ]:
```
members = clusters[clusters.merged_rep==cluster].member
```

In [ ]:
```
for m in members:
    print(m)
```

```
Q2RAL3
B7E663
Q7XJV3
Q94CS5
Q6ZBC3
Q8LJL3
Q10N90
Q8L4I2
Q5JL91
Q7EY69
Q6EPT2
A0A0N7KFK3
Q84TB9
A0A0P0XYR1
Q2QZ01
Q6YY31
Q7G2B2
Q6EPT4
Q6K9R5
A3ADD6
Q0JCK8
```

Looking at the `examples/AVRPik/AVRPik_vs_rice_clusters/merged_clusters/cluster_Q6K9R5.pdb_B/session.pse` file in PyMol, we can visualize the cluster and produce a figure like this:

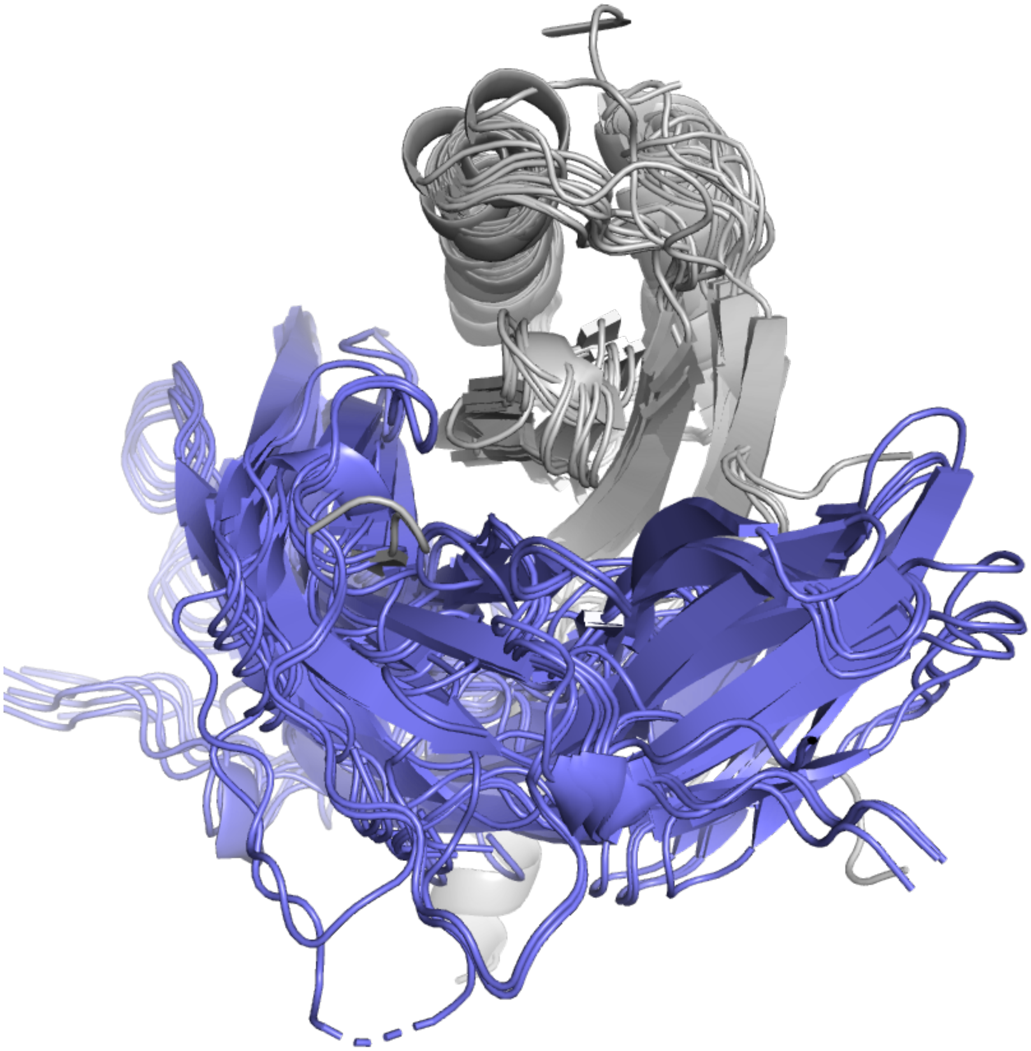

When aligning the HMA domains of the binder structures (grey), the different locations of the AVR-Pik protein raise the posibility of a more flexible binding across the entire surface of the HMA domain's beta strand.

Tip: In PyMol, you can remove the residues with low pLDDT score to make the figure cleaner, with the command:

```
select b < 50; remove sele
```
